# Temperate phages enhance host fitness via RNA-guided flagellar remodeling

**DOI:** 10.1101/2025.07.22.666180

**Authors:** Matt W.G. Walker, Egill Richard, Tanner Wiegand, Jing Wang, Zaofeng Yang, Americo A. Casas-Ciniglio, Florian T. Hoffmann, Hamna Shahnawaz, Ryan G. Gaudet, Nicholas Arpaia, Israel S. Fernández, Samuel H. Sternberg

## Abstract

Bacterial flagella drive motility and chemotaxis while also playing critical roles in host-pathogen interactions, as their oligomeric subunit, flagellin, is specifically recognized by the mammalian immune system and flagellotropic bacteriophages. We recently discovered a family of phage-encoded, RNA-guided transcription factors known as TldR that regulate flagellin expression. However, the biological significance for this regulation, particularly in the context of host fitness, remained unknown. By focusing on a human clinical Enterobacter isolate that encodes a Flagellin Remodeling prophage (FRφ), here we show that FRφ exploits the combined action of TldR and its flagellin isoform to dramatically alter the flagellar composition and phenotypic properties of its host. This transformation has striking biological consequences, enhancing bacterial motility and mammalian immune evasion, and structural studies by cryo-EM of host- and prophage-encoded filaments reveal distinct architectures underlying these physiological changes. Moreover, we find that FRφ improves colonization in the murine gut, illustrating the beneficial effect of prophage-mediated flagellar remodeling in a host-associated environment. Remarkably, flagellin-regulating TldR homologs emerged multiple times independently, further highlighting the strong selective pressures that drove evolution of RNA-guided flagellin control. Collectively, our results reveal how RNA-guided transcription factors emerged in a parallel evolutionary path to CRISPR-Cas and were co-opted by phages to remodel the flagellar apparatus and enhance host fitness.

## INTRODUCTION

Tight and dynamic regulation of gene expression is fundamental to cellular and organismal function, occurring at both the transcriptional and post-transcriptional levels^1,2^. Transcription factors (TFs) typically regulate gene expression by binding specific genomic regulatory regions through protein-DNA interactions^3^, while post-transcriptional control often relies on small RNAs that recognize target transcripts via RNA-RNA base pairing^4^. In contrast, we recently reported the widespread existence of programmable, RNA-guided TFs whose sequence specificity is conferred by RNA-DNA complementarity^5^. By exploring a family of transposon-encoded, RNA-guided nucleases within the TnpB superfamily — which are evolutionary precursors to CRISPR-Cas9 and Cas12^6,7^ — we uncovered multiple independent genesis events of TnpB nuclease domain inactivation, without an apparent loss of guide RNA (gRNA) and DNA binding activities. Intriguingly, many of these <u>T</u>npB-like nuclease-<u>d</u>ead repressors (TldRs) evolved strong genetic associations with novel non-transposon genes, indicative of molecular domestication and functional coupling^5,8^.

In one unusual TldR clade, we uncovered a novel isoform of flagellin (FliC) encoded adjacent to the TldR-gRNA cassette, and further identified both genes as residing within dormant, chromosomally integrated bacteriophages known as prophages^5^. Flagellin is the major extracellular structural component of the bacterial flagellum, a large membrane-embedded organelle that drives motility^9^. FliC is composed of a conserved D0/D1 core domain required for filament polymerization and motor function, flanked by highly variable, surface-exposed domains that diversify extensively across species^9^. The TldR-associated flagellin homologs represent the first known example of prophage-encoded flagellin (hereafter FliC_P_), and are typically highly divergent from their host-encoded counterparts (FliC_H_) (**Extended Data Fig. 1 a**). Remarkably, we revealed that in some clinical isolates of *Enterobacter*, prophage-encoded TldR specifically represses FliC_H_ expression by targeting its promoter^5^, allowing the FliC to outcompete FliC and thus transform the flagellar identity of *Enterobacter* (**Fig. 1a**).

**Fig. 1.**
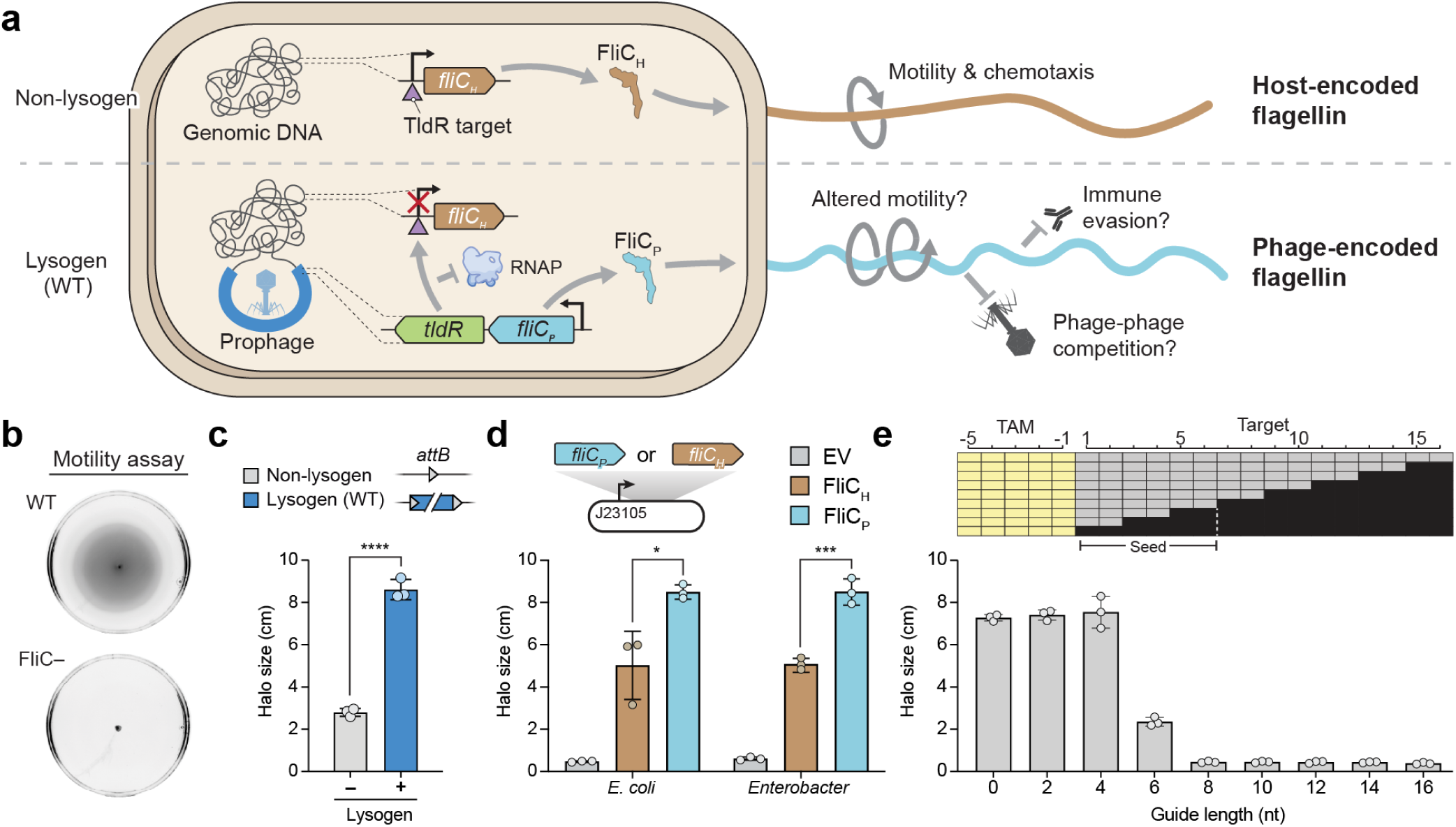
RNA-guided flagellin regulation in *Enterobacter* enhances cellular motility. **(a)** Schematic overview of TldR-mediated flagellar transformation. In the non-lysogenic state (top), the host-encoded flagellin gene *fliC_H_* is actively transcribed and assembled into a native flagellum (brown). In the lysogen (bottom), a prophage-encoded RNA-guided transcription factor (TldR) represses *fliC_H_* transcription, allowing expression of the prophage-encoded flagellin homolog *fliC_P_*, which assembles into a distinct flagellar filament (blue). This isoform switch has the potential to modulate bacterial motility, immune evasion, and susceptibility to secondary phage infection. **(b)** Representative soft agar plates imaged after 18 h of growth. The top image shows the wild-type lysogenic strain (WT), which exhibits a clear motility halo indicative of active flagellar function. The bottom panel shows a non-motile FliC– strain lacking both the host and prophage-encoded flagellin genes (Δ*fliC_H_ Δprophage*), resulting in complete loss of motility. **(c)** Bar graph quantifying bacterial motility via halo size measurements from experiments performed as in **B**, for the indicated strains. Bars indicate mean ± s.d. (n = 3 biological replicates); **** p < 0.0001 (unpaired t-test). **(d)** Bar graph quantifying bacterial motility in *E. coli* AW405 or *Enterobacter* sp. BIDMC93 lacking endogenous flagellin, with episomal expression of either FliC_H_ or FliC_P_; an empty vector (EV) serves as a negative control. Bars indicate mean ± s.d. (n = 3 biological replicates); * p < 0.05; *** p <0.001 (unpaired t-test). **(E)** Bar graph quantifying bacterial motility in *E. coli* AW405 expressing TldR along with a guide RNA targeting the *fliC* promoter region. The schematic above illustrates the positions of specific mutations (black rectangles) relative to an invariant cognate target adjacent motif (TAM). Mutations within the seed region (nucleotides 1–6) de-repress FliC and thus lead to an increase in observed motility, indicating its necessity for efficient RNA-guided gene silencing. Bars indicate mean ± s.d. (n = 3 biological replicates).

The integration of temperate phages into the bacterial chromosome — a process known as lysogenization — can lead to drastic phenotypic changes in the host, a phenomenon referred to as lysogenic conversion. In some cases, such as *Vibrio cholerae*^10^, lysogenic conversion has transformed non-pathogenic strains into potent human pathogens. Other prophage-conferred traits include enhanced biofilm formation^11^, improved cell invasion and intracellular survival^12^, antibiotic resistance^13^, resistance to phages^14^, and, more recently, competitive interactions with other bacteria^15^. However, rarely are lysogenic conversion events associated with alterations to host machinery as complex and fundamental as the flagellum, leading us to investigate the range and nature of this remodeling event.

In this study, we sought to understand how <u>F</u>lagellin <u>R</u>emodeling pro<u>ph</u>ages (FRφ) impact host fitness by transforming their flagellar composition. We reveal that rewiring of the host flagellar apparatus by FRφ significantly increases bacterial motility while decreasing recognition by the TLR5 receptor in human cells. Strikingly, FRφ also increases the ability of its host to colonize the murine gut, highlighting how phage-encoded organellar changes can impose substantial consequences on the physiological success of its host. Finally, we also uncovered an independent clade of TldR homologs that converged on the use of gRNAs to regulate flagellin isoform expression, highlighting the strong selective pressures driving flagellar diversification. These RNA-guided regulators are furthermore encoded adjacent to the translational regulator CsrA, providing insights into a broader paradigm of dual coordinated transcriptional and post-transcriptional gene regulation.

## RESULTS

### RNA-guided flagellin regulation in *Enterobacter* enhances cellular motility

We set out to understand the biological implications of RNA-guided flagellin regulation, focusing our initial efforts on the primary function of bacterial motility. Motility can be quantitatively measured by spotting bacteria onto a soft (0.2%) LB-agar plate, where motile strains form a visible halo from the inoculation point (**Fig. 1b**), thus serving as a proxy for locomotion, In our initial assays, we compared *Enterobacter* sp. BIDMC93 (hereafter *Ent*) cells with and without the prophage encoding *fliC_P_*. We used lambda Red-based recombineering to knock out the entire 53.5-kb prophage and reconstitute the *attB* site (**Materials and Methods**), and observed a striking motility enhancement in cells containing the prophage (**Fig. 1c**). To test the specific genetic factors encoded within the prophage required for this effect, we generated targeted disruptions within the prophage to knock out *fliC_P_* or *tldR*, and also reprogrammed the guide region to a non-targeting (NT) sequence that no longer repressed *fliC_H_*. Each of these mutations significantly occluded motility enhancement (**Extended Data Fig. 1b**), confirming that the enhanced motility phenotype was dependent on prophage regulation of the host flagellin gene.

To confirm that this regulatory system is active in its native context, we examined the expression profile of all prophage genes within the lysogenic strain. RNA-seq revealed that both *tldR* and *fliC_P_* were potently expressed (**Extended Data Fig. 1c**), supporting their active role in host modulation. Furthermore, ChIP-seq of TldR revealed native targeting to the *fliC_H_* locus (**Extended Data Fig. 1d**), and RIP-seq confirmed the TldR association with its cognate RNA (**Extended Data Fig. 1e**). These data establish that in the native lysogen, the prophage-encoded TldR is expressed, associates with its guide RNA, and targets the host flagellin gene for regulation.

Having established that the RNA-guided system is active in *Ent* and required for motility enhancement, we next asked whether the increased motility was due to differences in flagellin expression or distinct biophysical properties of the filaments. To isolate the contribution of each flagellin isoform, we episomally expressed either FliC_H_ or FliC_P_ under the same constitutive promoter, in either *E. coli* and *Enterobacter* cells lacking endogenous flagellin. In both strain backgrounds, we observed a pronounced motility increase when FliC_P_ was expressed as compared to FliC_H_ (**Fig. 1d**), indicating that the prophage-encoded flagellin filaments possess intrinsic properties that are responsible for the enhanced motility.

Next, we exploited the robust motility phenotype to better understand the molecular basis of TldR-mediated repression. As in other CRISPR systems that rely on RNA-guided targeting^6,16^, we hypothesized that the TldR regulatory activity would be governed by specific gRNA features, such as a seed sequence and minimal complementarity requirement. When we disrupted guide-target complementarity, we found that guide lengths as short as 6 nucleotides (nt) yielded strong levels of flagellin repression and corresponding reductions in motility (**Fig. 1e**), reminiscent of the seed sequence used by Cas9 and Cas12^17,18^ and consistent with the experimentally observed length of other TldR homologs^5^. We further defined gRNA requirements by testing a panel of 2-nt guide-target mismatches, which underscored the requirement of a 6-nt seed sequence (**Extended Data Fig. 2a**). Additionally, when we screened the guide-target pairings naturally present within TldR homologs from other organisms, we found that potent gene repression was retained despite a spectrum of mismatches distributed beyond the seed sequence (**Extended Data Fig. 2b**). These results reveal a precise molecular mechanism by which a prophage-encoded TldR silences host flagellin production to dramatically enhance bacterial motility.

### Architectural determinants of FliC_P_-dependent motility enhancement

Next, we leveraged cryo-EM to directly visualize differences between FliC_P_ and FliC_H_ filaments that could explain their distinct motility phenotypes. We purified flagellin filaments by expressing both flagellin isoforms in an *E. coli ΔfliC* knockout strain, mechanically shearing filaments from cells, and isolating filaments via centrifugation (**Methods**) (**Fig. 2a**). Both filament structures were determined using helical reconstruction in Relion5^19^, with the host FliC_H_ filament reconstructed with helical symmetry to an overall resolution of 3.7 Å and the prophage FliC_P_ filament reconstructed using a hybrid approach to 7 Å (**Fig. 2b,c, Extended Data Fig. 3, Methods**).

**Fig. 2.**
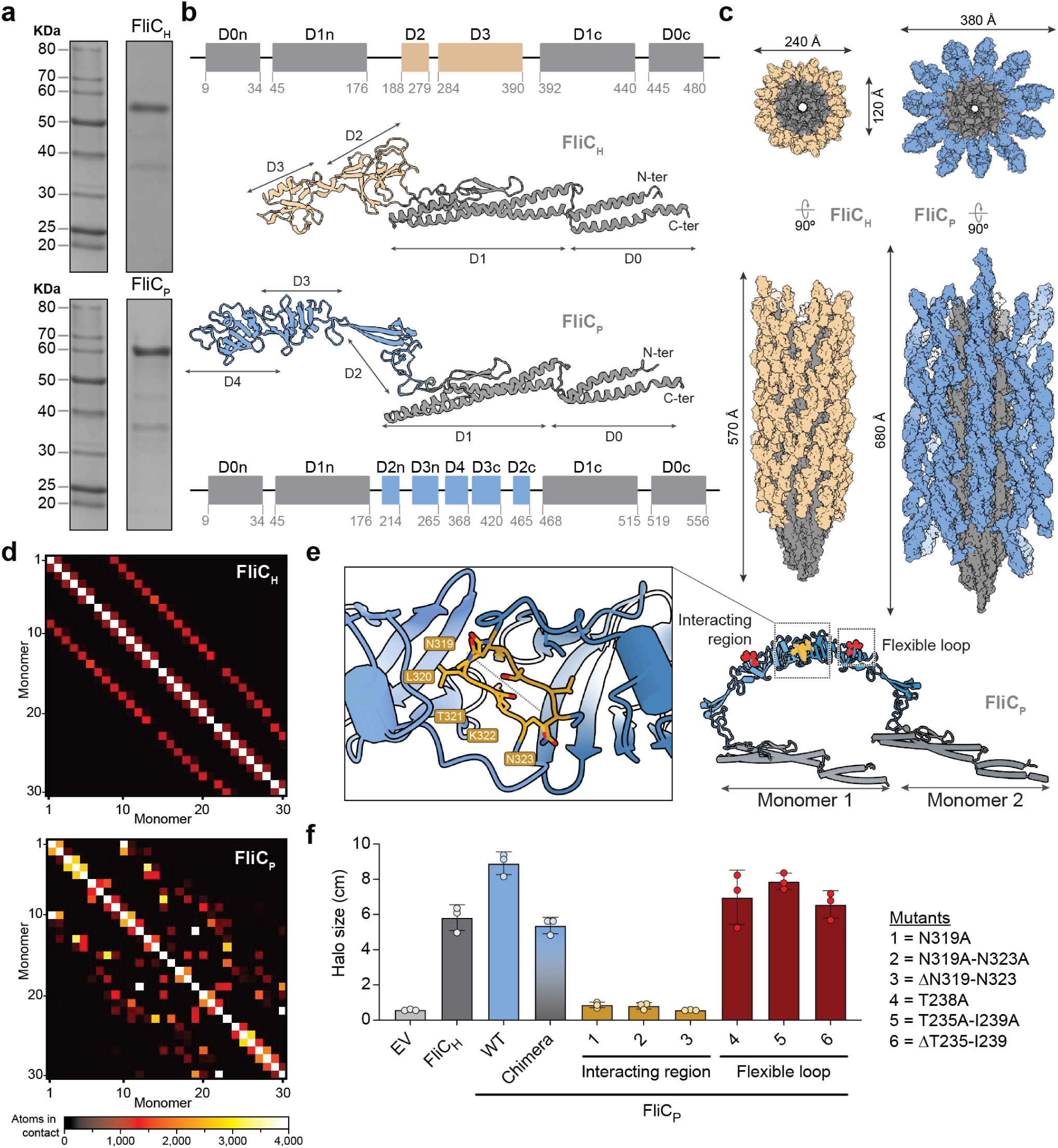
Cryo-EM structures of *Enterobacter* host- and prophage-encoded flagellin filaments. **(a)** Representative SDS-PAGE analysis of the flagellar fraction from deflagellated *E. coli* cells expressing recombinant *Ent*FliC_H_ (top) or *Ent*FliC_P_ (bottom), shown alongside a protein molecular weight ladder. **(b)** Domain annotations and cryo-EM structures of *Ent*FliC_H_ (top) and *Ent*FliC_P_ (bottom) monomers. Constant (D0–D1) and hypervariable (D2–D4) domains are labeled. **(c)** Molecular surface representation of *Ent*FliC_H_ (left) and *Ent*FliC_P_ (right) filaments from cryo-EM data, shown in both cross-sectional (top) and side (bottom) views. **(d)** Inter-subunit contact heatmaps for *Ent*FliC_H_ (top) and *Ent*FliC_P_ (bottom) filaments. Each heatmap quantifies the number of atomic contacts between thirty monomers within the respective filament, defined as any pair of atoms with distance ≤ 5 Å. This cutoff was selected to capture a broad range of non-covalent interactions, including van der Waals contacts, aromatic stacking, and proximal side chain interactions. **(e)** Cryo-EM structural model of two interacting FliC_P_ monomers (light and dark blue), highlighting the interaction region (inset, orange) and flexible loop (red) used for the experiments in panel (F). **(f)** Bar graph quantifying bacterial motility in an *Enterobacter* strain lacking endogenous flagellin, with episomal expression of FliC_H_, FliC_P_, or FliC_P_ variants harboring the indicated mutations. The empty vector (EV) serves as a negative control, and the chimeric construct substitutes the variable domain of FliC_H_ with that of FliC_P_. Bars indicate mean ± s.d. (n = 3 biological replicates).

While FliC_P_ and FliC_H_ share a conserved core composed of highly similar D0 and D1 domains, their external domains (D2–D4) differ markedly in composition (**Fig. 2b**). The N- and C-termini of each FliC_P_ and FliC_H_ monomer interact to form long alpha-helical bundles that contact neighboring protomers, assembling with a periodicity of 11 monomers per turn around a ∼25 Å pore that runs through the ∼120 Å-wide core and is used to export additional FliC copies during flagellar polymerization (**Fig. 2c**). Extending beyond these core regions, which bear strong similarity to flagellar filaments from *Escherichia coli* or *Salmonella enterica*^20^, variable domains project from the external surface of the filament, comprising the majority of the solvent exposed surface area (**Fig. 2c**). In FliC_H_ filaments, the external D2 and D3 domains form only minimal contacts with neighboring FliC_H_ subunits (**Fig. 2d**), creating threaded channels that run clockwise down the length of the ∼240 Å-wide filament (**Fig. 2c**). In contrast, the external variable domains of FliC_P_ (D2, D3, and D4) form loops that extend dramatically from the flagellar core, shielding the inner filament within a much larger (∼380 Å) reticulum of interacting subunits. FliC_P_ protomers form only minimal interactions with neighboring subunits through these variable domains, instead engaging in symmetric interfaces with subunits located roughly two helical turns apart (**Fig. 2d,e**). These extended loops impart flexibility to the mesh-like network surrounding the FliC_P_ core, in contrast to the more rigid contacts formed between variable domains of adjacent FliC_H_ monomers (**Extended Data Fig. 3e**). Overall, FliC_P_ variable domains mediate a substantially greater number of side chain contacts between subunits than observed in FliC_H_ filaments (**Fig. 2d**), which we suspect could FliC_P_ filament stability.

To test whether the FliC_P_ outer domain interactions contribute to enhanced motility, we designed a panel of mutations that disrupted these contacts. First, we substituted the entire FliC_P_ variable domain with the FliC_H_ variable domain and observed that the motility phenotype for this chimera perfectly phenocopied that of FliC_H_ (**Fig. 2f**). Next, we honed in on specific interacting residues within the FliC_P_ outer domain, focusing on a stretch of interacting residues between N319-N323 (**Fig. 2e**). Mutations within this region, including even a single residue substitution, caused a dramatic loss of cellular motility (**Fig. 2f**, variants 1-3). Mutating a distal stretch of residues outside the interaction interface did not impair motility (**Fig. 2f**, variants 4-6), lending support to the model that FliC_P_ interactions within the variable domain enhance motility.

### FRφ is an infectious temperate phage that produces *Siphoviridae*-like virions

Although our prior genomics analysis^5^ suggested that *tldR-fliC_P_*–containing prophages are likely to encode mobile viral agents, we nevertheless were not able to exclude the possibility that these were instead cryptic (inactive) prophages. Thus, we first asked whether the integrated prophage could enter the lytic phase and produce viral particles capable of infecting naive host cells. To detect novel lysogenization events, we employed a dual antibiotic resistance selection approach, in which we incorporated a kanamycin resistance cassette (*kanR*) into the prophage of a donor strain, and a chloramphenicol resistance cassette (*cmR*) into the genome of a *Δprophage* recipient strain (**Fig. 3a, Extended Data Fig. 4a**). Then, we treated the donor cells with mitomycin C (MMC), a DNA damaging agent typically used to induce prophages, filtered the treated donor cells to isolate phage particles <0.22 µm, and mixed the putative phage-containing supernatant with the *Δprophage* recipient cells. After plating cells on double-antibiotic media, we readily detected kanamycin- and chloramphenicol-resistant colonies, whereas no doubly-resistant colonies were observed when the assay was performed with control cells lacking the prophage (**Fig. 3b**). Intriguingly, we observed comparable levels of lysogenization even in the absence of MMC treatment (**Extended Data Fig. 4b**), indicating that the phage spontaneously produces ample infectious particles under standard liquid growth conditions. To confirm that newly isolated lysogens harbored integrated prophage genomes, we performed long-read whole genome sequencing and verified that the *kanR*-marked phage was integrated at the expected *attB* site (**Extended Data Fig. 4c**). To directly visualize the phage, we imaged the eluate by cryo-EM and observed phage particles with icosahedral heads and long, flexible, noncontractile tails — hallmarks of the *Siphoviridae* family^21^ (**Fig. 3c**). We thus designated this virus FRφ (<u>F</u>lagellin <u>R</u>emodeling phage), reflecting its unique regulatory function.

**Fig. 3.**
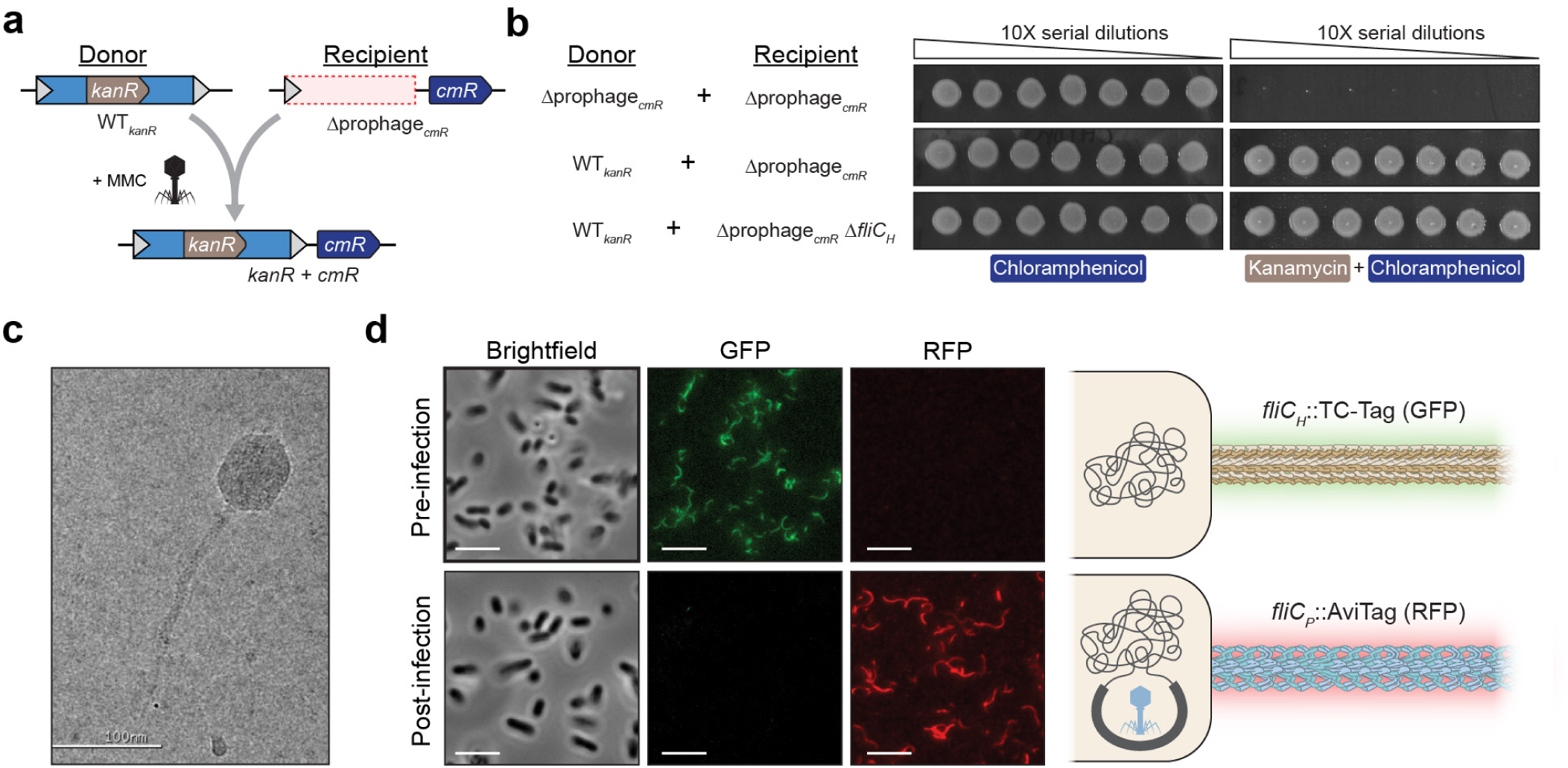
FRφ is a temperate, λ-like phage that modulates host flagellin composition. **(a)** Schematic of lysogenization assay, in which the prophage within the donor strain encodes a kanamycin resistance cassette (*kanR*, light green). The recipient strain lacks the prophage and encodes a chloramphenicol resistance cassette (*cmR*, blue) downstream of the attachment site (grey triangle), such that successful lysogenization events result in doubly-resistant clones. Prophage induction was induced with mitomycin C (MMC). **(b)** Representative plate images after the lysogenization assay in (A), demonstrating successful phage isolation and infection of naive recipient cells. Lysogenization efficiencies were indistinguishable when recipient cells lacked host flagellin (*ΔfliC_H_*). **(c)** Electron micrograph of FRφ. The scale bar represents 100 nm. **(d)** Representative microscopy images of cells before and after lysogenization, depicting the flagellar switch. Cells were co-labeled with FlAsH to detect TC-tagged FliC_H_, in the GFP channel, and streptavidin-Alexa Fluor 568 to detect AviTagged FliC_P_ in the RFP channel. The scale bar represents 5 µm.

Importantly, our lysogenization assay also allowed us to investigate whether host flagella are required for infection. Many phages employ superinfection exclusion mechanisms to prevent subsequent infection by the same or related phages^14^, and since some flagellotropic phages like χ use flagella as receptors^22^, we hypothesized that TldR-mediated flagellar transformation might serve as a superinfection exclusion strategy to prevent further infection by altering the flagellar composition of the infected cell. To test this, we infected a recipient strain lacking flagella (*Δprophage ΔfliC_H_*), and screened for doubly-resistant clones. Surprisingly, we observed no difference in the number of resulting colonies compared to infection experiments with a strain containing intact flagella (**Fig. 3b**), indicating that host flagellin is not required for infection and ruling out flagellar transformation as a mechanism of superinfection exclusion.

Motivated by the hypothesis that FRφ infection triggers a physical remodeling of the flagellum, we next performed fluorescence microscopy using isoform-specific labeling to visualize the switch from FliC_H_ to FliC_P_ in infected cells. One challenge in flagellin imaging is that direct fusion to traditional fluorophores such as GFP disrupts flagellin export and filament formation, and existing methods instead rely on small-molecule labeling techniques, such as cysteine-reactive maleimide dyes or the FlAsH system, which labels a genetically-encoded tetracysteine (TC) motif^23^. To complement these strategies with an orthogonal method, we pursued an *in vivo* biotinylation approach using the 15-amino acid AviTag, which can be efficiently biotinylated by the endogenous ligase BirA and detected with fluorescently labeled streptavidin^24^. We first established this method in *E. coli*, by scarlessly inserting the AviTag into *Eco*FliC at the same site previously used for tetracysteine tagging. Incubation with Streptavidin-Alexa Fluor 568 yielded robust flagellar signal even without exogenous biotin or BirA (**Extended Data Fig. 5a**), confirming efficient *in vivo* labeling. To verify that both labeling strategies were compatible, we co-labeled cells with FlAsH and the streptavidin conjugate and observed flagellin signal in the expected channels (**Extended Data Fig. 5b**). These *E. coli* benchmarking experiments confirmed the feasibility of multiplexed, isoform-specific labeling approaches.

We then applied this approach in *Enterobacter* to visualize flagellar transformation during FRφ infection. We scarlessly tagged *Enterobacter* FliC_H_ and FliC_P_ with the TC motif and AviTag, respectively, leveraging the atomic models from our cryo-EM structures to identify flexible regions and using a streamlined genome engineering method based on no-SCAR (**Methods, Extended Data Fig. 5c,d**). We then performed a lysogenization experiment, infecting *Δprophage* recipient cells expressing TC-tagged FliC_H_ with phage particles isolated from donor cells expressing AviTagged FliC_P_, and imaged cells before and after infection. This dual-labeling strategy revealed a striking and complete switch in filament composition from FliC_H_ to FliC_P_ (**Fig. 3d**), providing direct visual evidence of flagellar transformation upon phage infection.

### FRφ encodes a tail fiber protein shufflon locus that exploits FhuA for cellular entry

During long-read sequencing of our lysogenic *Ent* strain, we observed striking genetic heterogeneity within a predicted tail fiber protein (TFP) locus located upstream of the *fliC_P_*-*tldR* region (**Fig. 4a**). Closer inspection of the sequencing data revealed alternative 3′ coding sequences corresponding to three distinct C-terminal TFP isoforms (TFP-C1, -C2, and -C3). These differences appeared to arise from recombination events within a shufflon — a specialized genetic locus known to undergo site-specific rearrangements that diversity the C-terminal domains of phage or pilus-associated proteins. Quantification of isoform abundance using long-read sequencing confirmed that TFP-C1 was the dominant product (∼50% of reads), while TFP-C2 and TFP-C3 each accounted for ∼25% of the population (**Fig. 4b**). This distribution was consistent in our sequencing data of both the integrated prophage from *Ent* genomic DNA, as well as phage DNA from isolated virions, indicating that these isoforms are stably maintained across both chromosomal and viral contexts (**Fig. 4b**). Notably, the same heterogeneity was also detected in publicly available raw sequencing datasets for our strain, though it was not represented in the published isogenic genome, highlighting inherent limitations of reference genome assembly from genetically diverse, non-clonal bacterial populations. The presence of a TFP shufflon in FRφ suggested a mechanism for tail fiber diversification, yet the host receptor(s) for these isoforms remained unknown. Having established that FRφ infection does not require flagella (**Fig. 3b**), we next sought to identify which TFP variant(s) mediate infection, and what cellular receptor(s) they target.

**Fig. 4.**
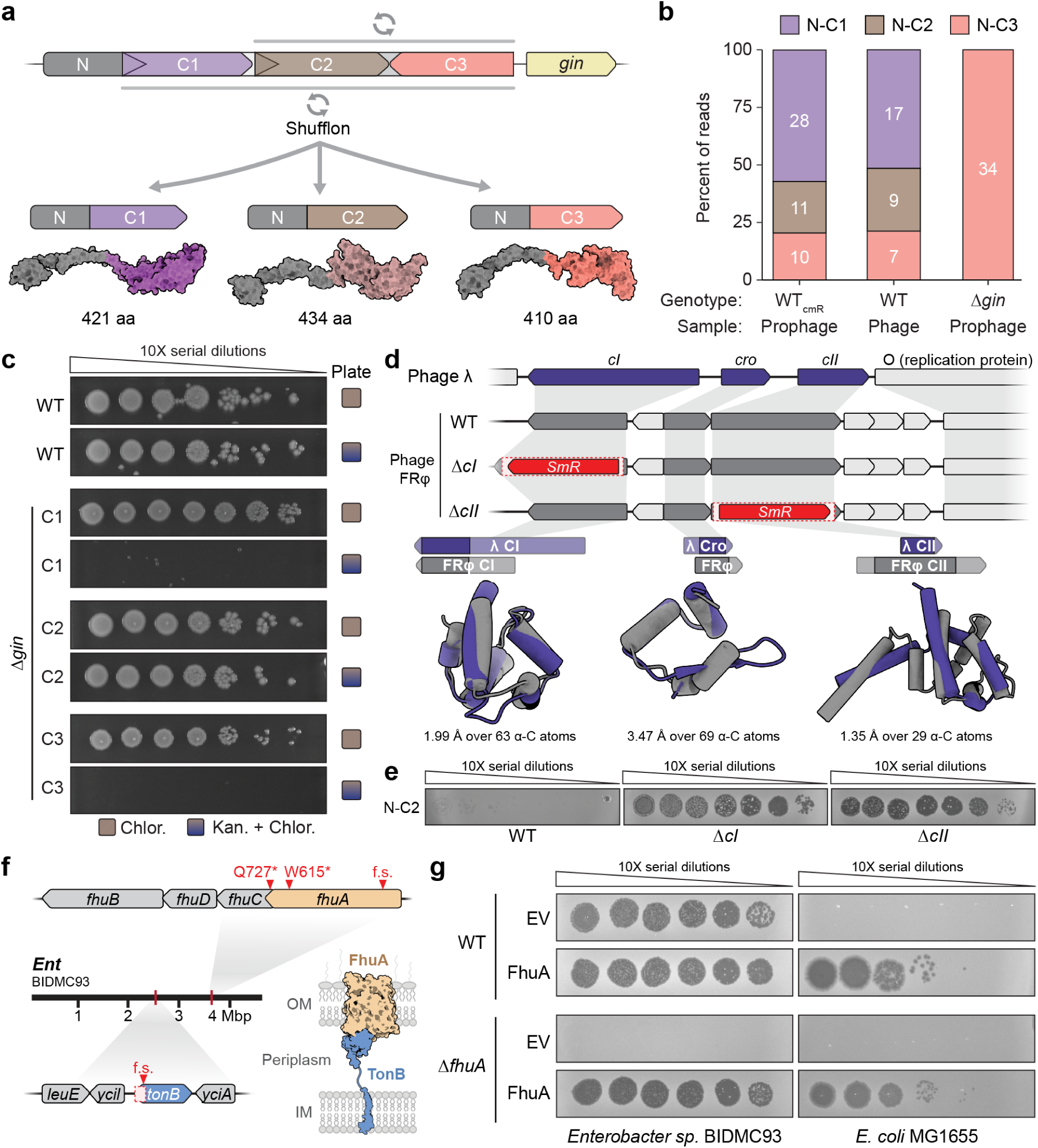
FRφ encodes a tail fiber protein shufflon locus that exploits FhuA for cellular entry. **(a)** Schematic of the FRφ-encoded shufflon tail fiber protein (TFP) locus, with AlphaFold 3 predictions of each TFP isoform. DNA segments capable of inversion are indicated at the top with circular arrows, and the distinct C-terminal domains are labeled C1–C3; *gin* encodes the responsible recombinase. **(b)** Quantification of the relative abundance of each TFP isoform from long-read sequencing of the WT FRφ prophage embedded in the *Ent* genome, isolated WT phage virions, and a prophage harboring a *Δgin* deletion mutation. Raw junction counts from HTS reads are listed in white text. **(c)** Representative plate images after lysogenization assay in a *Δgin* background, where TFP was locked into each of three distinct states harboring a unique C-terminal domain (C1, C2, or C3). Only phage particles expressing TFP-C2 can lysogenize *Ent*. **(d)** Comparison of phage λ lysogenic control locus and FRφ (top), and schematic depictions of *cI* / *cII* deletions to generate lytic/virulent FRφ (middle). Superpositions of homologous regions of CI, CII, and Cro are shown for phages λ and FRφ, depicted as AlphaFold 3 predictions, with RMSD values over *n* α-carbon atoms shown below each structural comparison; superpositions were calculated with the PDBeFold tool from EMBL-EBI. Regions covered by each superposition are indicated on the gene schematics above each predicted structure. **(e)** Plaquing assay of FRφ mutants *ΔcI* and *ΔcII* on a *Δprophage* strain of *Ent*, confirming that these mutations produce a lytic phage variant that generates clear zones of cell death. **(f)** Schematic illustrating the location and type of mutations in sequenced strains that are resistant to infection by FRφ. Three strains contain mutations in *fhuA*, while two contain mutations in *tonB*. **(g)** Plaquing assays demonstrate that FhuA complementation is necessary for FRφ infection in a *ΔfhuA* knockout strain, shown for both *Enterobacter* (left) and *E. coli* (right).

We returned to our lysogenization assay, but using a donor strain engineered to express a single, locked TFP variant. We began by deleting the *gin* recombinase located downstream of the TFP, which we hypothesized to be responsible for catalyzing shufflon rearrangements. Long-read sequencing of the *Δgin* donor strain confirmed the absence of recombination, resulting in a single, stable TFP isoform (**Fig. 4b**). Intriguingly, phylogenetic analysis of the *gin* gene revealed its homology to the Tn3-family resolvase TnpR (**Extended Data Fig. 6a, Table S1**) and uncovered related loci encoding up to six TFP paralogs in other bacterial genomes (**Extended Data Fig. 6b**), underscoring the broader evolutionary complexity of TFP diversification.

With donor strains locked into each of the three possible TFP states (C1, C2, or C3), we purified isogenic phage populations and repeated our lysogenization assay. Strikingly, only the TFP-C2 phage was capable of forming doubly-resistant colonies (**Fig. 4c**), identifying this variant as the sole isoform competent for *Ent* infection. This result not only revealed the functional specificity of TFP-C2 but also highlighted the potential for shufflon-mediated host-range modulation within the FRφ phage.

Next, to identify the host receptor used by FRφ, we decided to generate a lytic version of the phage and screen for phage-insensitive *Enterobacter* mutants. We began by deleting the putative *cI* and *cII* repressor genes, identified by homology to the well-characterized λ phage regulatory genes (**Fig. 4d**). Plaque assays confirmed that knockout of either *cI* or *cII* yielded virulent (non-temperate) forms of FRφ that produced clear zones of cell death indicative of productive infection and cellular lysis (**Fig. 4e**). We then applied a whole-plate plaquing strategy to isolate colonies that were resistant to lytic infection. Longread whole genome sequencing of five resistant clones revealed three putative loss-of-function mutations in *fhuA* and two in *tonB* (**Fig. 4f**). *fhuA* encodes a β-barrel outer membrane receptor previously implicated in phage entry for phages T1, T5, and phi80^25^, while *tonB* encodes a cytoplasmic membrane protein known to support the structural integrity and function of FhuA (**Fig. 4f**).

To validate that FhuA is required for FRφ infection, we engineered a Δ*fhuA* strain of *Ent* and observed a complete loss of plaque formation, supporting the hypothesis that FhuA functions as the essential receptor (**Fig. 4g**). To confirm this, we performed a rescue experiment by reintroducing an episomal copy of *fhuA* into the Δ*fhuA* strain, which restored phage infectivity (**Fig. 4g**). Remarkably, heterologous expression of *fhuA* in *E. coli* MG1655 — which is normally resistant to FRφ — was also sufficient to permit infection and lysis, demonstrating that FhuA alone is sufficient to confer susceptibility in a non-native host (**Fig. 4g**). Together, these findings identify FhuA as the key host receptor exploited by the TFP-C2 isoform of FRφ, linking shufflon-mediated tail fiber variation to host tropism and establishing a mechanistic basis for cellular entry.

### Screening for flagellotropic *Enterobacter* phage in environmental samples

Flagellotropic phages like χ use flagella as receptor^22^, which represents an attractive and large (μm-size) extracellular appendage. Given this observation, and the dramatically different structures of FliC_H_ and FliC_P_, we hypothesized that the flagellar transformation orchestrated by TldR might act as a potent strategy to endow lysogenic bacteria with novel antiphage resistance against competing lytic, flagellotropic phages. However, no *Enterobacter*-specific phages with flagellotropy have, to our knowledge, been identified.

We employed two distinct high-throughput screening methods aiming at isolating novel flagellotropic phages from wastewater, capable of infecting *Ent*. The first method involved isolating plaques obtained after mixing wastewater samples with a *Δprophage* strain expressing host FliC_H_, and then identifying flagellin-specific phages as those that, after enrichment, were unable to infect isogenic strains *lacking* FliC_H_. We isolated 632 phages through this first round of screening, but all of these phages were similarly capable of infecting the *Δprophage ΔfliC_H_ Ent* strain (**Extended Data Fig. 7a**). Hence, despite the high number of phages identified by this method, it did not reveal *Enterobacter*-specific flagellotropic phages.

Our second method was adapted from Phage DisCo^26^ and involved co-infection of a mixed population of strains expressing FliC_H_/RFP or FliC_P_/GFP. This approach immediately reveals phages that selectively infect one strain over the other via the formation of green or red ‘plaques,’ halving the infection experiments required to uncover FliC-specific infection differences. We successfully adapted this method to screen flagellotropic phages and carefully benchmarked it using *E. coli* and phages χ and T4 (**Extended Data Fig. 7b**), as well as in *Enterobacter* using our lytic FRφ phage (Δ*cI*) and *ΔfhuA* strains (**Extended Data Fig. 7c**). However, of the hundreds of phages screened using this method, we again failed to isolate flagellotropic phages (**Extended Data Fig. 7d**). In total, we screened over 1,100 phages but did not identify any with clear flagellotropic activity in *Enterobacter* (**Extended Data Fig. 7e**), suggesting that such phages are either rare, absent from our sampled environments, or require alternative screening strategies for detection.

### Flagellar remodeling promotes mammalian host immune evasion and improves gut engraftment

Having excluded superinfection exclusion and phage-phage competition as likely evolutionary driving forces behind flagellin remodeling, while nonetheless observing potent effects on bacterial motility, we next wanted to explore the impact on interactions with the mammalian innate immune system. Flagellin is detected as a key pathogen-associated molecular pattern (PAMP) by TLR5 extracellularly^27^ and NAIP5 intracellularly^28^, and some bacterial pathogens are known to regulate flagellin as a general strategy for enhancing colonization and pathogenesis. *Enterobacter* species from the *cloacae* complex, including *Enterobacter* sp. BIDMC93, are ubiquitous commensals in the human gastrointestinal tract but can become pathogenic in immunocompromised individuals. Since *Ent* is not an intracellular pathogen but would presumably be surveilled by the immune system via TLR5, we asked whether lysogenization by FRφ, and ensuing flagellar transformation, would alter TLR5 detection and downstream inflammatory signaling.

The epitope sequence that TLR5 binds^29^ is broadly conserved across FliC_P_ homologs, consistent with its location in the conserved D1 domain (**Extended Data Fig. 8**), but prior studies have demonstrated that this motif is not sufficient for TLR5 recognition and activation^30^. We therefore sought to test flagellin isoforms directly in a cellular model of TLR5 activation, using a human HEK293T cell line engineered to express TLR5 and a colorimetric reporter for NFκB-driven inflammation (**Fig. 5a**). When we stimulated HEK293T cells with the WT *Ent* lysogenic strain (expressing primarily FliC_P_) and compared it to a strain lacking any flagellin (*Δprophage* Δ*fliC_H_*), the WT triggered a robust TLR5-driven NF-κB activation signal, which we used as a baseline (**Fig. 5b**). Strikingly, a Δ*fliC_H_* strain expressing only the prophage-encoded FliC_P_ isoform elicited a similar signal, whereas a strain expressing only FliC_H_ (Δ*fliC_P_-tldR-gRNA*) triggered nearly twice the level of NF-κB activation. These data indicate that flagella composed of FliC_H_ are substantially more immunostimulatory than those composed of FliC_P_. To prove that this immune modulation is due to TldR-mediated silencing of *fliC_H_*, we examined NF-κB activation in strains expressing only TldR, with or without its guide RNA. Silencing of *fliC_H_* via TldR markedly reduced immune activation, while de-repression — either through deletion of the gRNA or substitution with a non-targeting (NT) sequence — restored the high NF-κB activation observed with FliC_H_ alone (**Fig. 5b**). Together, these results demonstrate that the FRφ phage diminishes flagellin immunogenicity by simultaneously encoding a less immunostimulatory flagellin isoform and a dedicated silencing system to repress the host copy.

**Fig. 5.**
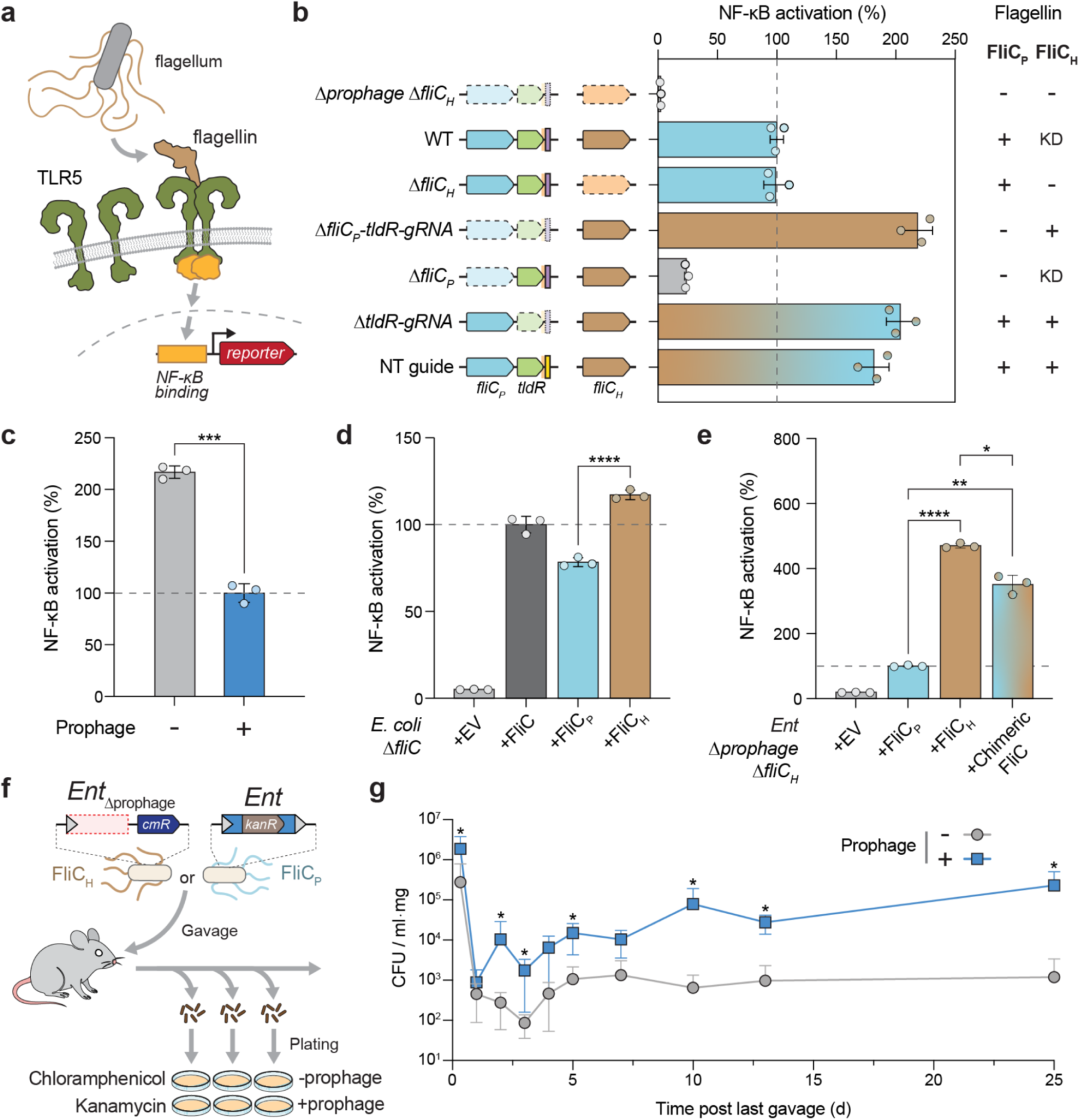
Phage-encoded flagellin attenuates activation of the mammalian host immune system and enhances engraftment. **(a)** Schematic of the TLR5 activation assay using human TLR5 expressing HEK293T reporter cells. The readout is a colorimetric assay measuring NF-κB activation. **(b)** Panel of *Ent* mutants and corresponding TLR5-driven NF-κB activation in HEK293T cells, expressed as a percentage of the signal obtained with the WT lysogenic strain. The flagellin isoform that is expressed is indicated at right: absent (–), expressed (+), knocked-down by TldR (KD). Bars indicate mean ± s.d. (n = 3 biological replicates). **(c)** Bar graph quantifying TLR5 activation by *Ent* with and without the FRφ prophage. Bars indicate mean ± s.d. (n = 3 biological replicates); *** p-value = 0.000131 (unpaired t-test) **(d)** Bar graph quantifying TLR5 activation by *E. coli* AW405 episomally expressing *E. coli* FliC, *Ent*FliC_H_ or *Ent*FliC_P_. Bars indicate mean ± s.d. (n = 3 biological replicates); **** p-value = 5.648e-05 (unpaired t-test). **(e)** Bar graph quantifying TLR5 activation by *Ent* episomally expressing FliC_H_, FLiC_P_ or a chimeric flagellin consisting of the conserved FliC_P_ domains (D0-D1) and variable FliC_H_ domains (D2-D3). Bars indicate mean ± s.d. (n = 3 biological replicates); **** p-value = 1.968e-05, ** p-value = 0.003951, * p-value = 0.01374 (unpaired t-tests). **(f)** Schematic of gut colonization assay, in which mice are gavaged with *Ent* strains containing or lacking the FRφ prophage. Each strain is marked with a unique antibiotic cassette, such that plating of bacteria from feces over time enables quantification of the colonization level. **(g)** Line graph showing results from the mouse gut colonization assay described in (F), quantified as colony-forming units (CFU) per mL·mg of resuspended feces for Ent strains with or without the prophage. Each data point represents the mean of five mice; error bars indicate standard deviation. Statistical significance was assessed using a Mann–Whitney test with FDR correction (q < 0.05); asterisks denote significant differences.

We next deleted the entire prophage and found that the non-lysogenic strain exhibited substantially higher levels of TLR5-driven NF-κB activation (**Fig. 5c**), consistent with our targeted genetic perturbations of *fliC_H_* and *fliC_P_*. To further confirm that these differences could be specifically ascribed to flagellin, we recombinantly expressed FliC_H_ or FliC_P_ under carefully controlled, episomal plasmid conditions, in *ΔfliC* strains of either *E. coli* or *Enterobacter*. Even in these distinct heterologous contexts, FliC_P_ consistently elicited a weaker NF-κB activation signal than FliC_H_ (**Fig. 5d,e**), supporting the interpretation that this differential effect is a property of flagellin and/or the flagellar filament it comprises. Intriguingly, a chimeric FliC comprising the conserved domains (D0-1) of FliC_P_ and variable domain (D2-3) of FliC_H_ largely phenocopied FliC_H_ (**Fig. 5e**), strongly suggesting that the solvent-exposed outer domain of the flagellar filament, rather than the TLR5 recognition motif itself, dictates the levels of TLR5 binding and/ or signaling.

Finally, we investigated how flagellar transformation and the ensuing effects on motility and TLR5 activation might impact the ability of *Enterobacter* to colonize the gut, using the mouse as a model. Mice were gavaged with isogenic *Ent* strains either lacking or carrying the FRφ prophage, in which the integrase gene was deleted (Δ*int*, see **Methods**) to prevent excision during the course of the colonization assay. Colonization was then monitored over 25 days by plating feces and quantifying abundances via drug marker selection (**Fig. 5f**). Remarkably, the lysogenic strain expressing FliC_P_ achieved significantly higher fecal abundances than the non-lysogenic strain expressing FliC_H_, at all time points after 24 hours post-gavage until the end of data collection (**Fig. 5g**). Control experiments confirmed the absence of pre-existing, antibiotic-resistant strains in the gut microbiota (**Extended Data Fig. 9a**), and that gavage efficiencies were equivalent across groups, with ∼5×10^9^ CFU administered per dose over three consecutive days (**Extended Data Fig. 9b**). Importantly, increased persistence of the lysogen was not associated with signs of pathogenicity, as both groups of mice gained weight steadily over the course of the experiment (**Extended Data Fig. 9c**).

Collectively, these results highlight multiple axes of host-pathogen interactions that are manipulated through flagellar transformation by phage FRφ, with the infected bacterial host itself becoming a more competitive constituent of the microbiome within its own mammalian host. This bacterial fitness advantage likely served as one of the driving forces behind evolutionary emergence of the phage-encoded *tldR-fliC_P_* pathway for RNA-guided flagellin control.

### Dual modes of flagellin regulation by CsrA-associated TldRs

Are FRφ-family phages unique in regulating flagellin via RNA-guided transcription factors, given the far-reaching physiological impact of flagellar diversification? Intriguingly, our previous discovery of TldR function also reported a distinct clade of homologs genetically associated with both *fliC* and the post-transcriptional regulator *csrA*^11^. These loci were not phage-encoded, but instead chromosomally integrated within commensal bacterial species — primarily from the human gut microbiome — raising the possibility that these host-encoded systems employ a dual mode of flagellin regulation at the level of transcription and translation (**Fig. 6a-c, Table S2**). To investigate this hypothesis, we heterologously expressed several *tldR* loci in *E. coli* and used RIP-seq to identify TldR-associated gRNAs, which could not be predicted bioinformatically (**Fig. 6d, Extended Data Fig. 10a, Table S3).** Parallel ChIP-seq experiments, followed by *de novo* motif analysis from enriched peaks, strikingly revealed that the guide region used for targeting was again complementary to predicted target sites found in distinct, non-*tldR*-associated flagellin genes (**Extended Data Fig. 10b-e**). This observation strongly suggests that these host-encoded TldR homologs similarly target alternative flagellin genes for repression, but by blocking transcription elongation rather than transcription initiation, given the positioning of target sites within the *fliC* open reading frame and not at the promoter (compare **Fig. 1a and Extended Data Fig. 10d,e**). Next, we performed RIP-seq of FLAG-tagged CsrA in cells also heterologously expressing candidate *fliC* target mRNA 5′ untranslated regions (UTRs), with the goal of identifying potential CsrA ligands that could explain its function. Remarkably, we observed strong enrichment ∼50-nt upstream of the *fliC* start codon, of the very same target gene putatively repressed by TldR (**Fig. 6e**). Secondary structure predictions of this region revealed a canonical CsrA binding motif within a stem-loop structure (**Fig. 6f**), supporting a model in which CsrA binds directly to the *fliC* 5′ UTR to repress translation, consistent with previous reports on its function^31^. To assess the specificity of this interaction, we ranked all regions in the CsrA IP sample with coverage above 100 counts per million (CPM). Among these, only two regions showed greater than 3-fold enrichment relative to input: the *fliC* UTR (38.9-fold) and *csrB* (13.9-fold) (**Extended Data Fig. 10f**). *csrB* is a well-characterized small RNA that contains multiple high-affinity CsrA binding sites and acts as a molecular sponge to titrate CsrA from mRNA targets. The strong and specific enrichment of the *fliC* UTR highlights it as a direct, high-priority regulatory target of CsrA.

**Fig. 6.**
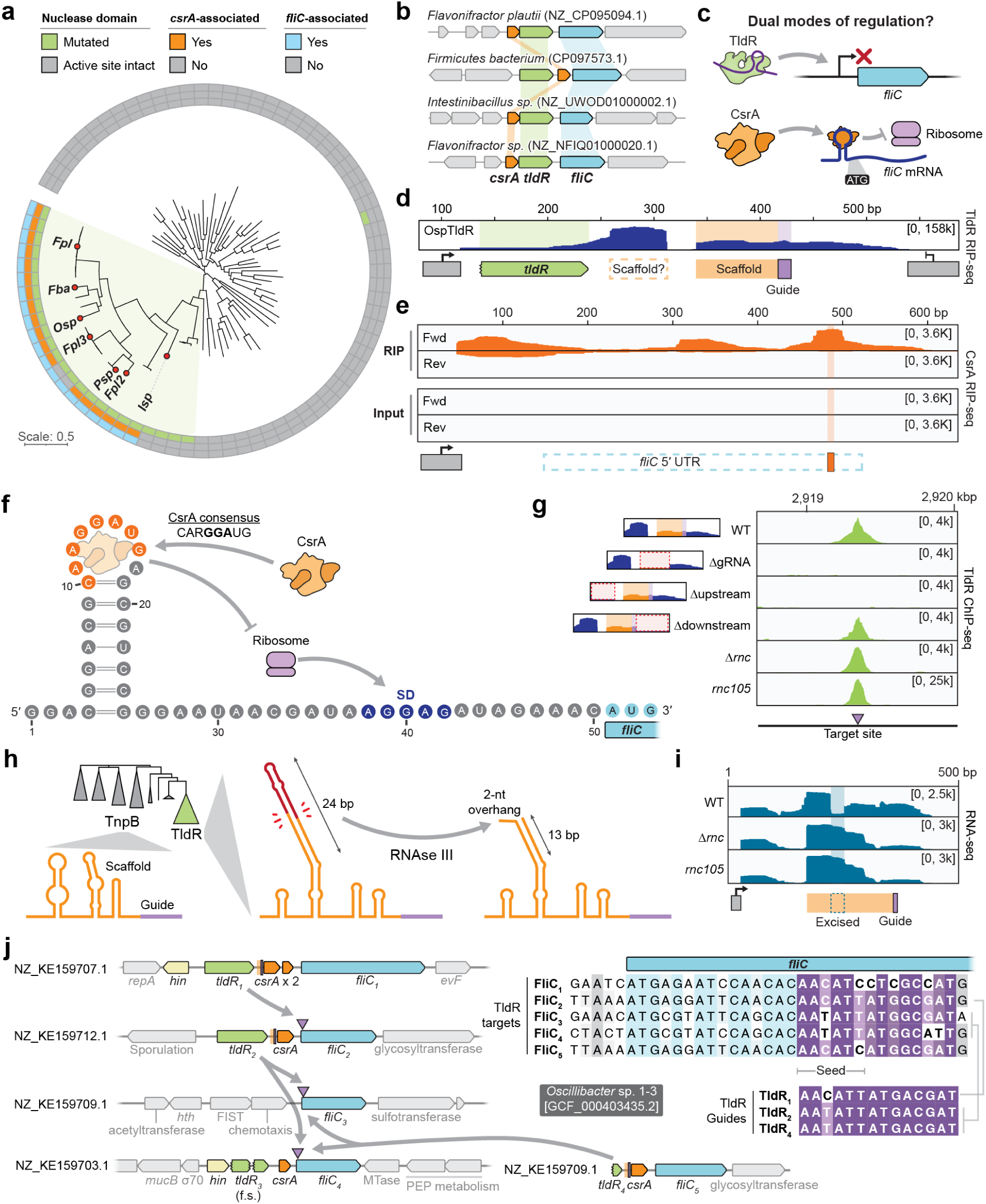
Dual modes of flagellin regulation by CsrA-associated TldRs. **(a)** Phylogenetic tree of CsrA-associated TldRs. The inner ring highlights TldRs with mutations in the RuvC nuclease domain (green); the middle ring indicates association with *csrA* (orange); the outer ring indicates association with *fliC* (also known as *hag* in *Firmicutes*). Red circles mark systems that were selected for expression and functional testing in *E. coli*. **(b)** Schematic of the genetic association and architecture of four *csrA*-associated *tldR* loci. **(c)** Model for dual modes of flagellin regulation. TldR targets the *fliC* gene to inhibit transcription, while CsrA targets the 5′ UTR of the *fliC* transcript to facilitate post-transcriptional repression. **(d)** RIP-seq coverage profiles of FLAG-tagged *Osp*TldR in *E. coli,* with the putative guide sequence highlighted in purple. Two regions of enrichment downstream of *tldR* are observed, corresponding to the predicted scaffold region of the guide RNA (orange). **(e)** RIP-seq coverage profiles of FLAG-tagged *Osp*CsrA in *E. coli*, mapped to the 5′ UTR of *fliC*. The orange highlight marks the predicted CsrA recognition motif (5′-CAAGGAUG-3′), where *Osp*CsrA is enriched on the forward strand in the IP sample. **(f)** RNA secondary structure prediction of the *fliC* 5′ UTR, showing the CsrA consensus sequence (orange) positioned within the loop of a predicted stem-loop structure. The Shine-Dalgarno (SD) and start codon sequences are highlighted in dark and light blue, respectively. **(g)** ChIP-seq coverage profiles of FLAG-tagged *Osp*TldR in *E. coli* at a genomic target site complementary to the guide sequence, shown across a panel of mutant backgrounds. Deletion of the guide RNA (ΔgRNA) or its upstream region (Δupstream) abolishes targeting, while deletion of the downstream region (Δdownstream) or inactivation of RNase III (*Δrnc* and *rnc105*) has no effect on TldR enrichment at the target site. **(h)** Predicted guide RNA structures from representative TnpB and TldR homologs, with a summarized version of the phylogenetic tree from (A). A schematic model of TldR gRNA before and after RNase III cleavage (right), shows activity that was facilitated by the expansion of a stem loop in the scaffold sequence. **(i)** RIP-seq coverage profiles of FLAG-tagged *Osp*TldR in *E. coli* across the guide RNA, comparing WT, *Δrnc*, and *rnc105* strains (top). Inactivation of RNase III restores coverage within the central dropout region, consistent with RNase III-dependent processing. **(j)** Schematic of the genetic architecture of four distinct *tldR-csrA* loci that are present in *Oscillibacter*, suggesting a complex and multi-tiered level of *fliC* regulation to control flagellin isoform expression. One locus contains frameshift (f.s.) mutations in the *tldR* gene, suggesting it is inactive, though it may still contribute gRNAs *in trans*. Contig names are shown next to each locus where they are encoded, and putative TldR regulatory targets and gRNA guide sequences are also shown (right).

In the course of analyzing the RIP-seq data for a *Oscillibacter* TldR homolog, we unexpectedly observed two peaks in the gRNA region separated by a severe drop in read coverage (**Fig. 6d**), in contrast to the continuous gRNA coverage observed for other TnpB and TldR homologs^5,32^. Subsequent perturbation experiments revealed that the upstream sequence was essential for DNA target binding, as assessed by ChIP-seq, whereas the region downstream of the guide sequence itself was dispensable (**Fig. 6g**). When we inspected the predicted gRNA secondary structure in greater detail, we noticed that the gap in coverage corresponded precisely with the distal end of a long stem-loop that would be present in the precursor gRNA transcript, leaving a characteristic 2-nt 3′ overhang reminiscent of RNase III cleavage products^33^ (**Fig. 6h**). Indeed, when we performed RNA-seq in *E. coli* strains harboring two distinct RNase III (*rnc*) loss-of-function mutations^34^, we found that continuous coverage across the full-length gRNA transcript was completely restored, without any loss of RNA-guided DNA binding activity (**Fig. 6g,i**). These results reveal a naturally occurring, RNase III-dependent dual-guide RNA that arises from a simple extension of a stem-loop feature present in the ancestral TnpB-associated gRNA, suggesting a potential evolutionary intermediate in the emergence of dual-guide RNAs (crRNA-tracrRNA) found in Type II and Type V CRISPR-Cas systems^35,36^.

In addition to probing gRNA maturation, we were curious to bioinformatically investigate the likely flagellar transformation executed by TldR. Interestingly, we found that the representative *Oscillibacter* strain encodes four distinct *csrA-tldR* loci, and careful inspection of each gRNA guide sequence identified a remarkably complex regulatory web of predicted cross-system flagellin repression (**Fig. 6j**). This genomic architecture suggests that TldR-CsrA systems likely act in a multiplexed fashion, forming a network that fine-tunes the expression of multiple *fliC* homologs across the genome. The co-occurrence of multiple transcriptional and translational regulatory nodes within a single strain underscores the overall complexity of these systems, further highlighting the selective pressures driving flagellar diversification via RNA-guided control in both prophage and host contexts.

In a final conceptual parallel with the flagellar remodeling activity of FRφ in *Enterobacter*, we also observed *hin* genes encoded immediately upstream of most *csrA*-*tldR* loci. Hin invertases are homologous to the Gin recombinases that swap bacteriophage tail fiber protein genes^37^ (**Fig. 4a**), leading us to search for local structural rearrangements, and indeed, comparative genomic analyses of closely related *csrA-tldR* loci from *Flavonifractor plautii* revealed ∼120-bp DNA inversions immediately downstream of *hin*, in a region containing a FliA-dependent promoter (**Extended Data Fig. 10g**). In *Salmonella*, Hin-mediated recombination of a DNA region encoding the flagellin gene promoter (i.e., an inverton) drives phase variation by toggling between expression of two flagellin isoforms^38^. The conserved positioning of *hin* and DNA inversions upstream of *csrA–tldR* loci supports a similar model, suggesting that a third layer of regulatory control modulates flagellar isoform expression combinatorially in organisms encoding TldR (**Extended Data Fig. 10h**).

## DISCUSSION

TnpB proteins encoded within bacterial IS*200*/IS*605* and IS*607* elements represent a vast reservoir of RNA-guided nucleases^6–8^ that have been independently domesticated numerous times over evolutionary timescales, leading to the emergence of dozens of unique CRISPR–Cas12 subtypes, with roles in bacterial adaptive immunity^39^ and programmable transposition^40,41^. In parallel, mutational inactivation of the RuvC domain resulted in the evolution of TnpB-like nuclease dead repressors (TldR) that function as RNA-guided transcription factors^5^, akin to the CRISPR interference (CRISPRi) technologies engineered by humans^42^. In this study, we focused on Flagellin Remodelling prophages (FRφ) that natively encode both a FliC_H_-targeting TldR and an alternative flagellin isoform, FliC_P_, effectively rewiring the flagellar machinery of their lysogenized hosts. Using an integrative experimental approach in *Enterobacter*, we demonstrate that FRφ-driven flagellar recomposition confers enhanced motility and decreased recognition by the innate immune receptor TLR5, ultimately increasing the capacity of *Enterobacter* to colonize the murine gut. This discovery highlights a major physiological change driven via lysogenic conversion, with potent ramifications for both the bacterial host as well as its mammalian host.

We initially envisioned that flagellar reprogramming by FRφ might serve as part of a superinfection exclusion (SIE) strategy, by which prophages protect their lysogenic host from subsequent infection by competing phages. This process has been extensively described for other phages such as T5, which expresses the lipoprotein Llp to block FhuA after infection, which is recognized as the receptor by both T5 and other phages^43^. More recently, Maxwell and Davidson and colleagues demonstrated that *Pseudomonas* prophages can alter the composition of the type IV pilus of their host to provide resistance against competing phages^44,45^. Within this context, one might hypothesize FRφ could bind FliC_H_-derived flagella as a receptor during infection, and then mask the receptor after lysogenic conversion by switching expression to the prophage-encoded FliC_P_ isoform; such a mechanism would protect the lysogen from reinfection-induced stress and reduce the likelihood of prophage induction into the lytic cycle, thereby stabilizing the integrated FRφ. Instead, our data revealed that FRφ, like T5 and ɸ80, recognizes FhuA as an extracellular receptor. Intriguingly, the combinatorial tail fiber protein isoforms catalyzed by the shufflon cassette likely increase the FRφ host range by allowing recognition of multiple receptors. We also considered the possibility that FRφ-mediated flagellar remodeling confers resistance to other flagellotropic phages, but failed to isolate any despite screening over 1,100 phage isolates from diverse environments. While we cannot exclude their existence, these data suggest such phages are rare and unlikely to be a dominant selective pressure. Thus, we conclude that flagellar recomposition in *Ent* is unlikely to have evolved primarily for phage resistance — either against secondary FRφ infections or against unrelated flagellotropic phages.

We next sought to understand what selective advantages might arise from FRφ-mediated lysogeny beyond phage exclusion. Strikingly, we found that *Enterobacter* strains featuring prophage-derived flagella exhibited markedly enhanced motility. This effect was recapitulated when FliC_P_ was heterologously expressed in *E. coli*, indicating that the phenotype is intrinsic to the flagellin isoform itself, rather than the host background. Structural analysis revealed that FliC_P_ assembles into a uniquely thick flagellar filament characterized by enlarged outer domains that wrap the filament core in a mesh-like configuration. These domains form increased inter-protofilament contacts that appear central to the motility phenotype, since targeted mutagenesis of residues at these interfaces diminished motility in *Enterobacter* cells expressing FliC_P_ variants.

In addition to promoting motility, FRφ lysogeny confers a second, distinct advantage: reduced activation of host innate immunity via TLR5. Despite the fact that the TLR5 epitope lies within the conserved D1 domain shared by both FliC_H_ and FliC_P_, we observed that FliC_H_ expression consistently elicited approximately twice the level of TLR5 activation, compared to FliC_P_. This difference was independent of host factors, as it was also recapitulated during heterologous *E. coli* expression, and thus again reflects a property of the filament itself. Chimeric constructs in which the outer domains of FliC_P_ were replaced by those of FliC_H_ restored high TLR5 activation, suggesting that filament architecture — rather than primary sequence variation at the TLR5-binding site — is the key determinant of immune visibility. We hypothesize that the increased thickness of the FliC_P_ filament, as well as the enhanced inter-protofilament interactions, could account for both the improved motility and reduced TLR5 activation, as greater filament integrity likely limits flagellin shedding and thus the availability of monomeric substrates for immune recognition. This interpretation aligns with theoretical models of bacterial motility, which predict that filament thickness and stiffness are critical determinants of propulsion dynamics^46^, and with the observation that some bacteria exploit specific proteases to degrade extracellular FliC monomers, thereby evading immune detection^47^. In the case of FRφ, however, the strategy would be structural rather than enzymatic, with filament stabilization indirectly reducing monomer availability and blunting host immune recognition.

The phenotypic effects of lysogenization result in a consequential ecological outcome: enhanced colonization of the murine gut by *Enterobacter* strains lysogenized by FRφ. Given that both motility and innate immune evasion are critical for successful host colonization^48,49^, it is plausible that both FliC_P_-driven traits contribute to this enhanced fitness, although disentangling their relative contributions remains a challenge. Remarkably, we also discovered evidence for RNA-guided flagellar regulation in other key members of the human gut microbiome, but driven by host-encoded TldR-FliC loci that furthermore collaborate with CsrA, likely enabling a dual mode of both transcriptional and post-transcriptional gene regulation. The complex network of flagellin isoform diversification, with up to five distinct homologs being differentially regulated, resonates with prior descriptions of flagellin phase variation in gut-associated bacteria^50^. The ability to modulate the constant flagellar inner core with novel solvent-exposed variable domains may enhance motility through high-viscosity mucus or environments with varying osmolarity and pH^51^. Indeed, the presence of six distinct flagellin genes in *Vibrio cholerae* is thought to confer motility advantages under different physicochemical conditions^52^, and *Salmonella enterica* exploits flagellin phase variation for increased virulence and enhanced host epithelial surface interactions to promote gut colonization^38^.

While TldR-FliC_P_ expression appears constitutive under the laboratory conditions tested in our study, we cannot exclude the possibility that expression varies *in vivo*. If so, FRφ lysogeny may provide not only structural flagellar remodeling, but also a mechanism for phase variation, enabling dynamic adaptation of motility and immune evasion in a tissue-specific or temporally controlled manner. Such a model would position FRφ as a modulator of host–microbe interactions, optimizing the fitness of its lysogenic host in complex and fluctuating environments.

## LIMITATIONS OF THE STUDY

Our analysis of flagellin expression was performed under standard laboratory conditions, in which we observed that both FliC_P_ and the TldR-gRNA system are constitutively expressed. In this context, we conclude that the prophage mediates constitutive flagellar reprogramming, leading to a durable switch from FliC_H_-to-FliC_P_ flagella. However, we cannot exclude the possibility that this expression pattern is altered in more complex environments such as the mammalian gut. Indeed, it is plausible that gut-associated *Enterobacter* dynamically alternates between expressing both flagellin isoforms in response to environmental cues. We also note that our motility assays demonstrating improved swimming with FliC_P_ raise the question of why the host-encoded FliC_H_ is retained, since superior motility might be expected to ultimately lead FliC_P_ isoforms to be rapidly selected by the host and fixed in the population. The fact that this is not observed by our comparative genomics analyses suggests the likelihood that FliC_H_ offers other advantages in ecological contexts not captured in our laboratory assays. Future work will be needed to understand the selective pressures shaping the maintenance of both flagellin variants in natural settings.

## DATA AVAILABILITY

High-throughput sequencing data have been deposited at the National Center for Biotechnology Information (NCBI) Sequence Read Archive (BioProject Accession: PRJNA1256936) and are publicly available.

## CODE AVAILABILITY

All original code has been deposited at Zenodo and is publicly available at DOI 10.5281/zenodo.15307114.

## ACKNOWLEDGMENTS

We thank I. Lisevich, R. Colin, and V. Sourjik for their careful reading of the manuscript. We are grateful to M. Baym and E.A. Rand for discussions related to Phage DisCo, and for providing wastewater samples from Boston. We also thank K.A. Human and the Bozeman Water Reclamation Facility for generously providing wastewater samples. We thank E.C. Greene and V.B. Raina for access to Amersham Typhoon imaging systems, and L.F. Landweber, J.E. Dworkin, M.A. Hydorn, and S.N. Nagarajan for access to fluorescence microscopy instrumentation. We thank M.E. Edwards and I.I. Ivaylo for assistance with anaerobic bacterial culturing, and D.R. Gelsinger for beat-beating training. We thank C. Meers for identifying CsrA-TldR homologs and providing bioinformatics support, and G.D. Lampe for assistance with mammalian cell culture. We thank members of the S.H. Sternberg lab for helpful discussions, T. Smith for general laboratory support, and the staff at the JP Sulzberger Columbia Genome Center for NGS support.

M.W.G.W. was supported by a National Science Foundation Graduate Research Fellowship (DGE-2036197). R.G.G. was supported by the National Institutes of Health (NIH DP2AI177904) and the Kinship Foundation through the Searle Scholar Program. N.A. was supported by the NIH (R01CA259634). I.S.F. was supported by the Spain Ministry of Science Innovation and Universities Grant (PID2023-147463NB-I00). Cryo-EM sample preparation and data collection were performed at the Simons Electron Microscopy Center at the New York Structural Biology Center, with support from the Simons Foundation (SF349247) and NIH (U24GM129539). S.H.S. was supported by the NSF Faculty Early Career Development Program Award (CAREER 2239685), a Pew Biomedical Scholarship, an Irma T. Hirschl Career Scientist Award, the Howard Hughes Medical Institute Investigator Program, and a generous startup package from the Columbia University Irving Medical Center Dean’s Office and the Vagelos Precision Medicine Fund.

## AUTHOR CONTRIBUTIONS

M.W.G.W., E.R., T.W., and S.H.S. conceived the project. M.W.G.W. performed strain engineering, motility assays, flagellin purifications, lysogenization assays, lytic infection assays, fluorescence microscopy, phage DisCo, and TldR-CsrA assays. E.R. performed strain engineering, lysogenization assays, lytic infection assays, phage hunting, TLR5 immunology experiments, and mouse gut colonization experiments. T.W. performed all bioinformatics analyses and assisted in data interpretation and visualization. J.W. prepared cryo-EM samples and collected cryo-EM data. Z.Y. performed mouse gut colonization experiments. A.A.C.-C. cloned flagellin expression vectors and performed motility and plaque assays. F.T.H. performed CsrA-TldR ChIP-seq experiments. H.S. performed and assisted in the design of TLR5 immunology experiments. R.G.G. advised on the design and interpretation of TLR5 immunology experiments. N.A. advised on the design and interpretation of mouse gut colonization experiments. I.S.F. advised on the design of cryo-EM experiments and determined all cryo-EM structures. S.H.S. oversaw the project. M.W.G.W., E.R., T.W., I.S.F., and S.H.S. discussed the data and wrote the manuscript, with input from all authors.

## METHODS

### Strain and plasmid construction

Strains used in this study are listed **Table S4**. Genomic mutants of *E. coli* and *Enterobacter* were generated either by Lambda Red recombineering^53^ or by an adapted protocol for scarless genomic edits using counterselection by *Sp*Cas9 (no-SCAR^54^). For strains constructed using Lambda Red recombineering, mutants were designed to replace each gene of interest with an antibiotic resistance cassette (either kanamycin, chloramphenicol, or spectinomycin), which was amplified by PCR with Q5 High-Fidelity DNA Polymerase (NEB) using primers that contained at least 50-bp homology arms flanking the disrupted locus. PCR amplicons were resolved on a 1% agarose gel and purified (QIAGEN). Electrocompetent cells were prepared containing a temperature-sensitive plasmid that encodes the Lambda Red machinery under the control of a temperature-sensitive promoter (pSL2684). Protein expression from the temperature-sensitive promoter was induced by incubating cells at 42 °C for 25 min immediately prior to electrocompetent cell preparation. Then cells were transformed with 200-600 ng of each insert (2 kV, 200 Ω, 25 µF) and recovered for at least 4 h in 3 ml of LB media. After recovery, cells were spread onto 100 mm standard plates with antibiotic (either 50 µg/ml kanamycin, 25 µg/ml chloramphenicol, or 200 µg/ml spectinomycin) and grown at 37 °C. Antibiotic-resistant colonies were genotypes by Sanger sequencing (GENEWIZ) to confirm each desired disruption.

For *E. coli* and *Enterobacter* strains constructed using *Sp*Cas9, we followed the protocol for Scarless Cas9 Assisted Recombineering (no-SCAR^54^), with some modifications that we found to be necessary for constructing genomic mutants of *E. coli* strain AW405. Spacers targeting the disrupted loci of interest were selected and cloned into a gRNA expression plasmid (pSL6050). Cells for the strain of interest were transformed with the gRNA expression plasmid at 30 °C (since it is temperature sensitive), then grown up to mid-log phase and induced with L-arabinose to 0.2% for 15 min. Cells were then made electrocompetent by washing twice in 10% glycerol, and co-transformed with ssDNA donor and ∼250 ng Cas9 plasmid (pSL0411). The ssDNA donor was designed using 30-50-bp homology arms to the desired edit site, with mutations that were introduced to prevent Cas9 cleavage after the ssDNA donor was used for repair, and was synthesized as a 4 nmol Ultramer (IDT). All ssDNA oligos used for scarless recombineering are listed in **Table S5**. After transformation, cells were recovered at 30 °C for 2 h, before being plated on 25 µg/ml chloramphenicol, 200 µg/ml spectinomycin and 0.1 µg/ml anhydrotetracycline and incubated overnight. The next day, single colonies were checked by PCR and Sanger sequencing (Genewiz) to identify colonies with the desired edit. Then, the temperature sensitive gRNA expression plasmid was cured by growing cells at 42 °C. To cure the Cas9 plasmid, cells were made chemically competent and transformed with a Cas9-targeting sgRNA plasmid (pSL0407). After transformation, cells were recovered at 30 °C for 3 h, induced with anhydrotetracycline to 0.1 µg/ml, then continued recovery for an addition 2 h before plating on 200 µg/ml spectinomycin and 0.1 µg/ml anhydrotetracycline, and incubating at 30 °C overnight. The next day single colonies were grown up without antibiotics and the sgRNA plasmid was cured by overnight growth at 42 °C. Cultures were checked for curing of both Cas9 and sgRNA plasmid by spotting onto plates containing either chloramphenicol (25 µg/ml) or spectinomycin (200 µg/ml).

The original *Enterobacter sp.* BIDMC93 strain encodes three distinct flagellin genes: one located within the FRφ prophage (*fliC_P_*), one chromosomal copy that is targeted and silenced by TldR (*fliC_H_*), and a third, distinct chromosomal homolog that we refer to as *fliC_2_*. To assess the prevalence of this additional flagellin gene, we performed a systematic analysis of 396 genomic assemblies containing *fliC_P_–tldR*. Across these genomes, we identified 893 flagellin homologs. Our analysis revealed that an extra *fliC_2_*-like copy was present in only ∼4.4% of *fliC_P_–tldR*–containing genomes, whereas a TldR-targeted *fliC_H_*-like homolog co-occurred with *fliC_P_–tldR* in approximately 93% of cases. Based on these findings, we concluded that *fliC_2_* represents a rare and unregulated flagellin gene not integrated into the canonical TldR regulatory system. Therefore, to enable unambiguous functional analysis of FliC_P_ and FliC_H_, we deleted *fliC_2_* from the *Enterobacter sp.* BIDMC93 background (*Enterobacter sp.* BIDMC93 *ΔfliC_2_*) and used this newly generated strain in all subsequent assays (see **Table S4**).

Plasmids used in this study are listed in **Table S6**. Flagellin expression vectors were constructed by cloning flagellin genes under a medium-strength constitutive promoter (J23105) on a medium-copy pET vector with carbenicillin resistance. TldR and CsrA were cloned into expression vectors under the control of constitutive J23105 promoters, and guide RNAs were expressed using constitutive J23119 promoters. Derivatives of these expression plasmids were cloned using a combination of methods, including Gibson assembly, restriction digestion-ligation, and around-the-horn PCR. Plasmids were cloned, propagated in NEB Turbo cells, purified using Miniprep kits (QIAGEN), and verified by Sanger sequencing (Genewiz).

### Bacterial motility assays

Motility assays were performed using the soft agar method, essentially as previously described^55^. Overnight cultures were diluted 1:100 in LB supplemented with antibiotic, then grown with gentle shaking (∼120 rpm) to OD_600_ = 0.6 at 30 °C. Then, 2 µl of culture was pipetted onto the center of semisolid agar plates (2.5% Miller’s LB broth and 0.2% Bacto agar) and incubated at 30 °C before imaging. Images were captured on a Bio-Rad Gel Doc XR Imaging system using epi-illumination and automatic exposure. Halo size was measured by taking the average of the vertical and horizontal diameter measurements for three replicates.

### Flagellar filament purification

Flagellar filaments were purified by mechanical shearing and centrifugation, essentially as previously described^56^. Overnight cultures of each *E. coli* strain (with constitutive episomal flagellin expression) were grown in 100 µg/ml carbenicillin with gentle shaking (∼120 rpm) for 6 h. Cultures were then normalized by OD_600_ and centrifuged at 4,000 *g* for 5 min, then the pellet was resuspended in deflagellation buffer (1 M Tris-Cl (pH 6.5) and 100 mM NaCl). Flagellar filaments were mechanically sheared from cell bodies by passing cells through a 27-gauge needle 15-20 times, and the filament fraction was isolated from cell bodies by centrifugation at 10,000 *g* for 15 min. The supernatant containing flagellar filaments was removed and concentrated 1,000-fold through a 100-kDa filter (Amicon Ultra Centrifugal Filter). Then, the buffer was exchanged to motility buffer (0.5 mM CaCl_2_, 0.1 mM EDTA, 20 mM HEPES (pH 7.5)) via dialysis (Thermo Scientific Slide-A-Lyzer MINI Dialysis Unit).

### Cryogenic electron microscopy

3 µl of purified flagellin samples were applied to a glow-discharged holey carbon Quantifoil Au R1.2/1.3 grids (Electron Microscopy Science, Hatfield, PA), and blotted with filter paper using a FEI Vitrobot Mark III at 4 °C and 100% humidity (blot force = 0, and blot time = 3.0-4.5 s) then plunge-frozen into liquid ethane. Preliminary screening was performed in a Glacios instrument (ThermoFisher) to check for ice quality and sample concentration/quality. Grids showing good contrast and long and multiple filaments per hole were saved for high-resolution data collection on a Titan-Krios instrument equipped with a BioQuantum-K3 direct detector with an integrated energy filter. Movies were collected at a magnification of 105,000x corresponding to a pixel size of 0.844 Å. Exposure time and fluence were adjusted to ∼1e^−^/ pixel/frame and 60 frames per movie were collected with a defoci range of −0.8 to −2.2 µm for a total dose of 61.02 e^−^/Å^2^.

### Helical reconstruction and structure determination

Cryo-EM data processing was integrally performed in Relion5^19^ with ctf estimation in CTFFIND4^57^ via the Relion wrapper. Motion correction was performed in whole frame mode and particle picking was performed with a modified version of Topaz that allows automatic filament picking^58^. First, we selected 50 images of each dataset that were manually picked and extracted using canonical values of the *E. coli* flagellum structure as reference^56^ (*id est* 4.85 Å rise). Particle stacks were generated by sliding a box of 600 pixels 48.5 Å apart along the picked filament (10 times the canonical rise) and these particles were subsequently binned to a final size of 200 pixels. Two rounds of 2D classification (100 classes each, tau fuge parameter (T) 2 and 4) allowed the selection of 8 2D class averages showing secondary structure elements. This set of particles (∼3000) was used for training a Topaz^59^ model for automatic filament picking of the whole dataset that produced a large dataset of 3 million particles. Recursive 2D classifications with increasing T values (from 2 to 5) and selecting for the best looking 2D averages in terms of secondary structure detail and straightness rendered a final set of ∼125,000 particles. The Fourier transform of the final 2D averages revealed a pattern of layer lines reminiscent of previous studies of bacterial filaments by cryo-EM^56,58^. The presence of additional layer lines beyond the canonical helical core of the *E. coli* flagellum hinted towards multiple symmetries in the same filament. To explore this possibility, we created a cylindrical mask including the external domains of the filaments but excluding the canonical helical core formed by the coiled-coils encoded in the D0-D1 domains. This approach allowed us to eliminate (using the signal subtraction feature of Relion5^19^) the signal corresponding to the central helical core and reconstruct using helical reconstruction without symmetry, the signal corresponding to the external sheath using as initial reference a smooth cylinder^19,60^. This approach rendered low resolution (∼8 Å) but highly informative reconstructions whose interpretation was facilitated by the AlphaFold 3 (AF3) predictions of the monomer structures for *Enterobacter* FliC_H_ and FliC_P_. By manually docking of the AF3 prediction for the external domains of FliC_H_ in the low-resolution maps obtained by helical reconstruction without symmetrical impositions, we were able to recognize a unit of 5 consecutive monomers that seemed to be repeating along the filament. This observation prompted us to attempt a helical reconstruction of the whole filament using 5 times the canonical values of the *E. coli* central core, that is rise of 24.4 Å and twist of −33.5 ° (canonical values are rise 4.88 Å and twist 65.4 °). This refinement converged to a map of around 4 Å with correct features for a map of this resolution such as well defined secondary structure elements including β-strand separation and identifiable bulky side chains. Ctf-parameters refinement and estimation of local particle movement and signal filtering using the Relion5 bayesian polishing approach rendered a final map with overall resolution of 3.7 Å^61,62^. The final map for FliC_H_ shows excellent density for the canonical core including many side chains as well as unambiguous density for the external domains.

For the FliC_P_ case we applied a similar strategy, isolating via signal subtraction the external layer of the filament and performing a helical reconstruction without symmetry. The interpretation of this map revealed a nuanced pattern of interactions whose interpretation was possible using the AF3 prediction for the FliC_P_ monomer. The FliC_P_ monomer adopts two conformations that we termed “up” and “down” which differ in a close to 180 ° flip of the D2-D3-D4 domains relative to the canonical D0-D1 domains (similar to previously reported flagellum structures from soil bacteria^56^). One monomer in “down” conformation interacts with another monomer in “up” conformation located in an adjacent protofilament, creating a dimeric unit that is repeated. Along a vertical protofilament, the repetition pattern is “down-down-up” and “up-up-down” in the neighboring protofilament. However, this pattern is not compatible with the underlying canonical symmetry of the D0-D1 domains when considering the eleven protofilaments that constitute the entire flagellum. This inconsistency is solved by alteration of the “down-down-up” pattern introducing a symmetry disruption. To generate a map of the complete filament for the FliC_P_ we improved the resolution of the external sheltering layer by ctf-parameter refinement and bayesian polishing to ∼7 Å reconstructing helically but without imposing any symmetry. In parallel, we refined the helical core to ∼3.5 Å applying helical reconstruction and the canonical flagellum parameters (rise 4.88 Å and twist 65.4 °). We produced a composite map aligning the N-terminal helix of D1, visible in the external layer map and merging both maps to a final resolution of 7 Å.

For model building of FliC_H_, using UCSF Chimera^63^, we manually placed 88 monomers in the final unsharpened map, carefully adjusting the position of the D2-D3 external domains relative to the D0D1 canonical domains for every monomer. We then divided the fitted monomers into vertically aligned groups of 11 protofilaments and every proto-filament was stereochemically refined in real-space against the post-processed map using phenix-real-space refinement with secondary structure restraints activated^64^. Following this step, we performed a reciprocal-space refinement for every protofilament using REFMAC5^65^ with secondary structure restraints calculated in ProSMART^66^.

For FliC_P_ we employed a similar model-building approach but due to the limited resolution we restricted our model refinement to a single step of real-space refinement in phenix using tight secondary structure restraints to avoid overfitting of the atomic model.

### Flagellin imaging

TC-tagged flagella were labeled using FlAsH, essentially as previously described^67^, with some modifications. This protocol was adapted to label AviTag flagella using streptavidin-Alexa Fluor. Overnight cultures were diluted 1:100 in LB supplemented with antibiotic, and grown with gentle shaking (∼120 rpm) for 6 h. Cells were then labeled by incubating them with the appropriate dye in LB. For labeling TC-tagged flagella, cells were labeled with 2.5 µM FlAsH (Thermo Fisher) and incubated at 30 °C for 2 h. For labeling AviTagged flagella, cells were labeled with 10 µg/ml streptavidin-Alexa Fluor 568 conjugate (Invitrogen) and incubated at 30 °C for 2 h. For co-labeling cells, both 2.5 µM FlAsH and 10 µg/ml streptavidin-Alexa Fluor 568 were added to cells.

Microscopy tunnel slides were prepared using two pieces of double-sided tape between microscope slides (Fisherbrand, 75 x 25 mm) and a coverslip (Fisherbrand, 22 mm, No 1.5). The tunnel was treated with 25 µl of 0.1% poly-L-lysine (Sigma) for 5 mins, then washed with M9 media. 20 µl of labeled cells were pipetted into the tunnel and allowed to sit for 10 mins, to adhere to the coverslip, before being washed twice with 50 µl M9 to remove unadhered cells. Images were captured using a Zeiss Axiovert 200 inverted microscope equipped with a 100X oil objective and an Axiocam 506 monochrome camera. Images were pseudo-colored using ImageJ^68^ when merging two channels.

### Chromatin-immunoprecipitation followed by next generation sequencing (ChIP-seq)

ChIP-seq was performed essentially as described^69^, with some modifications. Cells were scraped from solid agar plates and resuspended in 40 ml of LB media. Cells were fixed by adding formaldehyde to final concentration of 1%, and nutated for 20 min at room temperature. Formaldehyde was quenched by adding 4.6 ml of 2.5 M glycine and nutating for 10 min. Cells were then centrifuged at 4000 *g* at 4 °C for 8 min, and the pellet was washed twice in cold 1× TBS. To normalize cell density across cultures, the equivalent of 40 ml of OD_600_=0.6 was transferred to a tube, and pelleted again by centrifugation. The pellet was then transferred to a 1.5 ml Eppendorf tube and centrifuged at 10,000 *g* at 4 °C for 5 min. The supernatant was discarded and pellets were flash-frozen in liquid nitrogen.

Frozen cell pellets were resuspended in lysis buffer (50 mM HEPES-KOH (pH 7.5), 0.1% (w/v) sodium deoxycholate, 0.1% (w/v) SDS, 1 mM EDTA, 1% (v/v) Triton X-100, 150mM NaCl, 1× cOmplete Roche protease inhibitor), and transferred to a 1 ml milliTUBE AFA Fiber (Covaris). Cells were sonicated on an M220 Focused-ultrasonicator (Covaris) with the following SonoLab 7.2 settings: minimum temperature, 4 °C; set point, 6 °C; maximum temperature, 8 °C; peak power, 75.0; duty factor, 10; cycles/ bursts, 200; sonication time, 17.5 min. After sonication, 10 µl of cleared lysate was withdrawn and kept as the pre-IP control (“input”). The remainder was used for immunoprecipitation.

For immunoprecipitation, 25 µl Dynabeads Protein G (Thermo Fisher Scientific) for each sample was washed four times in 1× PBS + 0.5% BSA, then mixed with 4 µl monoclonal anti-FLAG M2 antibody produced in mouse (Sigma-Aldrich F1804), which was conjugated at 4 °C for 6 h. After conjugation, beads were washed four times with 1× PBS + 0.5% BSA, resuspended in 30 µl lysis buffer, then mixed with sonicated samples and rotated at 4 °C overnight.

The following day, beads were washed with lysis buffer lacking protease inhibitor three times, then washed once with lysis buffer lacking protease inhibitor and supplemented with 500 mM NaCl, once with ChIP wash buffer (10 mM Tris–HCl (pH 8.0), 250 mM LiCl, 0.5% (w/v) sodium deoxycholate, 0.5% (v/v) Nonidet-P40, 1 mM EDTA) and finally twice with TE buffer (10 mM Tris-HCl (pH 8.0), 1 mM EDTA). After the final wash, beads were resuspended in 200 µl elution buffer (1% (w/v) SDS, 0.1 M NaHCO_3_) and incubated at 65 °C for 1 h 15 min to release protein-DNA complexes from the beads, vortexing every 15 min to resuspend the beads. While incubating the immunoprecipitated samples, 10 µl of the frozen pre-IP samples were withdrawn and 190 µl elution buffer was added. After the 65 °C incubation was complete, 10 µl of 5 M NaCl was added to each pre-IP and IP sample, which were incubated at 65 °C overnight to reverse crosslinks.

After the overnight incubation, RNA digestion was performed using 1 µl RNase A (Thermo Fisher Scientific) and incubating at 37 °C for 1 h. Protein digestion was performed using 2.8 µl of 20 mg/ ml proteinase K (Thermo Fisher Scientific) and incubating samples at 55 °C for 1 h. DNA was purified (QIAquick PCR Purification Kit) and eluted in 40 µl TE buffer before the concentration was measured by fluorometry (DeNovix dsDNA Ultra High Sensitivity Kit). Samples were normalized to the lowest concentration, and DNA was prepared for next-generation sequencing using the NEBNext Ultra II DNA Library Prep Kit for Illumina (NEB). Samples were sequenced using an AVITI Sequencing System with a 2×150 Cloudbreak Freestyle Kit (Element Biosciences).

After sequencing, paired-end reads were filtered and trimmed using fastp^70^ and mapped to the *E. coli* MG1655 or *Ent* reference genome using bowtie2^71^. Then, we used Samtools to sort, index, and filter multi-mapping reads^72^. Finally, coverage was normalized to counts per million (CPM) using the deepTools2 command bamCoverage^73^ (v3.5.1).

### RNA immunoprecipitation (RIP-seq) of RNA bound by TldR or CsrA

Cells harvested for RIP-seq were cultured as described for ChIP-seq using an *E. coli* MG1655 strain. Cells were scraped from solid agar plates and resuspended in 1 ml of TBS buffer (20 mM Tris-HCl pH 7.5, 0.15 M NaCl). Next, cell density was measured (OD_600_) and an equivalent of 20 ml of OD_600_=0.5 was withdrawn into a new tube. Cells were pelleted by centrifugation at 4,000 *g* and 4 °C for 5 min, the supernatant was discarded, and pellets were flash frozen in liquid nitrogen.

Cell pellets were resuspended in 1.2 ml RIP lysis buffer supplemented with cOmplete Protease Inhibitor Cocktail (Roche) and SUPERase•In RNase Inhibitor (Thermo Fisher Scientific). Cells were then sonicated for 1.5 min total (2 sec ON, 5 sec OFF) at 20% amplitude. Lysates were centrifuged for 15 min at 4 °C at 21,000 *g* and the supernatant was transferred to a new tube. At this point, a small volume of each sample (24 μl, or 2%) was set aside as the input material.

For immunoprecipitation, 60 μl Dynabeads Protein G (Thermo Fisher Scientific) were first washed 3 times in 1 ml RIP lysis buffer (20 mM Tris-HCl pH 7.5, 150 mM KCl, 1 mM MgCl2, 0.2% Triton X-100), then resuspended in 1 ml RIP lysis buffer and combined with 20 μl anti-FLAG M2 antibody (Sigma-Aldrich). Samples were rotated at 4 °C for at least 3 h. Antibody-bead complexes were washed 3 times and resuspended in 60 μl RIP lysis buffer, then mixed with sonicated samples and rotated at 4 °C overnight.

The following day, samples were washed three times with ice-cold RIP wash buffer (20 mM Tris-HCl, 150 mM KCl, 1 mM MgCl_2_). After the final wash, beads were resuspended in 1 ml TRIzol (ThermoFisher Scientific) and RNA was eluted from the beads by incubating at RT for 5 min. The supernatant was separated using a magnetic rack and transferred to a new tube, then combined with 200 μl chloroform. Each sample was mixed vigorously by inversion, incubated at RT for 3 min, and centrifuged for 15 min at 4 °C at 12,000 *g*. RNA was isolated from the upper aqueous phase using an RNA Clean & Concentrator kit (Zymo). RNA from input samples was isolated in the same manner, using TRIzol and column purification. High-throughput sequencing library preparation was per-formed as described below for total RNA-seq of Enterobacter strains. Libraries were sequenced on an AVITI Sequencing System with a 2×150 Cloudbreak Freestyle Kit (Element Biosciences).

Adapter trimming, quality trimming, and read length filtering of RIP-seq reads was performed as described below for total RNA-seq experiments. Trimmed and filtered reads were mapped to the *E. coli* MG1655 reference genome using bwa-mem2v2.2.1^74^, with default parameters. Mapped reads were sorted, indexed, and converted into coverage tracks as described below for total RNA-seq experiments.

### Total RNA sequencing of *Enterobacter*

For RNA-sequencing on *E. coli* WT and the two RNase III knockout strains (*Δrnc* and *rnc105*), colonies were first scraped from solid media, resuspended in LB media, and the equivalent of 1 mL of OD_600_=1 culture was withdrawn. Samples were centrifuged at 3,000 *g* at 4 °C for 5 min, then the supernatant was discarded. RNA was extracted by adding 750 µl TRIzol, incubating for 5 min at room temperature, then 150 µl chloroform was added to each tube, shaken vigorously to mix, and left for 5 min at room temperature. Samples were centrifuged at 12,000 *g* at 4 °C for 15 min, then 400 µl of the upper aqueous layer was carefully transferred to a new tube. RNA was purified using an RNA Clean and Concentrator Kit (Zymo), eluting in 15 µl of DNase/RNase-free water. Samples were then diluted in NEBuffer 2 (NEB) to a total volume of 15 µl, and fragmented by incubating at 92 °C for 2 min. The fragmented RNA was simultaneously treated with RppH (NEB) and TURBO DNase (Thermo Fisher Scientific) in the presence of SUPERase•In RNase Inhibitor (Thermo Fisher Scientific) at 37 °C for 30 min, in order to remove DNA and 5′-pyrophosphate. For end repair, the RNA was treated with T4 PNK (NEB) in 1×T4 DNA ligase buffer (NEB), and incubated at 37 °C for 30 min. Samples were column-purified using an RNA Clean & Concentrator Kit (Zymo), and the concentration was determined using the DeNovix RNA Assay (DeNovix). Illumina adapter ligation and cDNA synthesis were performed using the NEBNext Small RNA Library Prep kit, using 100 ng of RNA per sample. High-throughput sequencing was performed on an AVITI Sequencing System with a 2×150 Cloudbreak Freestyle Kit (Element Biosciences).

RNA-seq reads were processed using cutadapt^75^ (v4.2) to remove adapter sequences, trim low-quality ends from reads, and exclude reads shorter than 15 bp. Trimmed and filtered reads were aligned to the MG1655 reference genome using bwa-mem2^74^ (v2.2.1) in paired-end mode with default parameters. SAMtools^72^ (v1.17) was used to filter for uniquely mapping reads using a MAPQ score threshold of 1, and to sort and index the unique reads. Coverage tracks were generated using bamCoverage^73^ (v3.5.1) with a bin size of 1, read extension to fragment size, and normalization by counts per million mapped reads (CPM) with exact scaling. Coverage tracks were visualized using IGV^76^.

### *Enterobacter* phage induction and isolation

Phage induction was performed using mitomycin C treatment, essentially as previously described^77^ with minor modifications. Overnight cultures of *Enterobacter* sp. BIDMC93 (*Ent*) strains were diluted 1:100 in LB media, and incubated at 37 °C with agitation until OD_600_=0.20. Then, cultures were treated with final concentration of 0.5 µg/ml mitomycin C and incubated at 37 °C for 16 h. Chloroform was added to a final concentration of 4% (v/v) and briefly vortexed to facilitate bacterial lysis, after which the lysate was centrifuged at 4000×*g* for 10 min to pellet cell debris. The supernatant was carefully withdrawn, taking care not to disrupt the chloroform-cell interface, and passed through a 0.22 µm filter. The phage-containing filtrate was stored at 4 °C.

### Lysogenization assay by dual-antibiotic resistance screening

To detect lysogenization, we used a custom assay that relied on labeling phage and bacteria with separate antibiotics and detecting cells that had acquired resistance to both antibiotics after mixing cells with phage particles. We used Lambda Red recombineering (as described above) to label the prophage with a chloramphenicol resistance cassette and the *Enterobacter* genome with a kanamycin resistance cassette. After inducing cells with mitomycin C and isolating phage-containing filtrate by sterile filtration (described above), 450 µl of recipient cells at OD_600_=0.20 were mixed with 50 µl of isolated phage particles, then incubated with shaking at 37 °C for 2 h before plating. Cells were serially diluted (10×) and plated as 4 µl spots onto LB agar supplemented with either single antibiotics (50 µg/ml kanamycin or 25 µg/ ml chloramphenicol) or double antibiotic plates (50 µg/ml kanamycin plus 25 µg/ml chloramphenicol). Plates were incubated at 37 °C for 16 h to allow the growth of colonies. Plates were imaged on a BioRad Gel Doc XR+ imager.

### Small drop plaque assays

Plaque assays were performed as previously described^78^. An overnight culture for each strain was mixed with 4 ml of freshly prepared molten soft agar (0.5% Difco agar + 2.5 % LB Miller’s broth) and poured over solid bottom agar (1.5 % agar + 2.5 % LB Miller’s broth, supplemented with the appropriate antibiotic) and rotated to evenly dispense the soft agar solution. The soft agar was left to solidify for 15 min at RT, during which 10× serial dilutions of phage samples were prepared in LB media. For plating, 3 µl of each phage dilution was spotted onto the surface of the soft agar, and plates were left to dry for 10 min under a laminar flow hood. Plates were incubated at 37 °C for 12-16 h to allow the formation of plaques. Plates were imaged on a BioRad Gel Doc XR+ imager.

### Isolation of FRφ-resistant mutants

*Enterobacter* cells with mutations in the FRφ receptor were isolated by performing whole-genome sequencing cells that grew after infection with lytic phage, essentially as previously described^79^. Overnight cultures of *Enterobacter Δprophage* cells were diluted 1:100 in LB media supplemented with 25 µg/ ml chloramphenicol and incubated at 37 °C until OD_600_=0.3. 1 ml of bacterial culture was mixed with 1 ml of lytic FRφ (Δ*cI* or Δ*cII*, constructed using Lambda Red recombineering, as described above). This mixture was incubated at 37 °C for 10 min, then plated on solid LB agar plates supplemented with 25 µg/ ml chloramphenicol, which were left overnight at 37 °C.

The next day, individual colonies were grown in LB media and checked by plaque spot assay (described above) to confirm resistance to FRφ. To identify mutations conferring resistance, cultures were sent for long-read whole genome sequencing (performed by Plasmidsaurus using Oxford Nanopore Technology with custom analysis and annotation). Annotated genomes generated from long-read sequencing data were analyzed for single nucleotide polymorphisms (SNPs) using Parsnp2^80^. This analysis revealed mutations in FhuA or FhuB. Mutations in FhuA were confirmed by rescue experiments, in which *Enterobacter* FhuA and its native promoter was cloned onto a plasmid (pSL7573), and used for transformation into *Enterobacter ΔfhuA* cells. Plaquing assays (described above) performed using this strain confirmed that expression of FhuA rescued the ability of FRφ to infect and lyse cells.

### Screening wastewater for flagellotropic *Enterobacter* phage

We screened wastewater for flagellotropic Enterobacter phage using two alternative methods, referred to as replica-plaquing and using the previously described Phage DisCo approach. For the replica-plaquing approach, we obtained wastewater from Bozeman (Montana), New York City (New York) and Boston (Massachusetts). The viral fraction of each sample was obtained by treatment with 4% chloroform, vortexing, spinning and passing the aqueous phase through a 0.22 µm filter. 200 ml of the wastewater viral fractions obtained in Bozeman and New York were then concentrated 100-fold by passing them through a 50 kDa filter to a final volume of 2 ml. The Boston viral suspension and the concentrated Bozeman and New York City samples were then used to obtain plates on an *Enterobacter ΔFRφ* lawn. Briefly, 100 µL of each sample were mixed with 100 µL of an exponentially growing culture of *Enterobacter ΔFRφ* at OD ∼ 0.3, incubated at room temperature for 5 minutes, then mixed into 4 ml of freshly prepared molten soft agar supplemented with 5 mM MgCl. The suspension was briefly inverted three times to mix, then poured over solid bottom agar (1.5 % agar + 2.5 % LB Miller’s broth, supplemented with the appropriate antibiotic). The soft agar was left to solidify for 15 min at RT and the obtained double layer agar plates were placed in a 37 °C incubator overnight. The following day, putative plaques were isolated using a 200 µl pipette tip and resuspended in 200 µl magnesium saline buffer (100 mM NaCl, 8 mM MgSO4 and 50 mM Tris at pH 7.5). Plaques isolated in this way are very small (< 1 mm) and may be confused with debris or bubbles in the soft agar. Therefore, to confirm that they can in turn form a larger plaque on a bacterial lawn, 3 µL of each resuspended putative plaque was pipetted onto a double-layer agar plate prepared with the *Enterobacter ΔFRφ* strain. Of the 868 putative plaques, 632 were confirmed as forming real plaques in this way. The corresponding plaques were isolated in 96-well plates, and the phage suspensions were treated with chloroform in a spin plate, wrung out and 100 µl of the supernatant pipetted into a new plate for storage at 4°C. The screen then consisted in stamping the contents of each plate, using a 96-pin replicator, onto double-layer agar plates containing either *Enterobacter ΔFRφ* or *Enterobacter ΔFRφ ΔfliC_H_*, in order to identify plaques that would form on the former but not on the latter. Of all the pages tested in this way, no phage exhibited this characteristic.

Phage DisCo was performed essentially as previously described^26^, with minor modifications. An equimolar ratio of mid-log phase cells for *Enterobacter Δprophage* cells expressing RFP under a constitutive promoter (J23119) were mixed with Δ*fliC_H_ Δprophage* cells expressing GFP under a constitutive promoter (J23105). A total of 100 µl of mixed strains plus 100 µl of phage suspension isolated from environmental samples were mixed with 4 ml of freshly prepared molten soft agar (0.5% Difco agar + 2.5 % LB Miller’s broth). The suspension was briefly inverted three times to mix, then poured over solid bottom agar (1.5 % agar + 2.5 % LB Miller’s broth, supplemented with the appropriate antibiotic) and rotated to evenly dispense the molten agar solution. The soft agar was left to solidify for 15 min at RT before placed in a 37 °C incubator overnight. The following day, plates were imaged on Amersham Typhoon Biomolecular imager (Cytiva) using 488 nm and 635 nm laser intensities. Images were pseudo-colored using ImageJ^112^ when merging two channels.

### TLR5/NF-κB Assay in HEK293T-Blue-hTLR5 cells

HEK293T-Blue-hTLR5 are HEK293T cells co transfected with the human TLR5 gene and an inducible SEAP (secreted embryonic alkaline phosphatase) reporter (Invivogen). The SEAP gene is placed under the control of the IFN-β minimal promoter fused to five NF-κB and AP-1-binding sites. Stimulation with a TLR5 ligand (i.e. flagellin) activates NF-κB and AP-1 which induces the production of SEAP. This enzyme induces a color shift from pink to blue of the chromogenic substrate in the HEK-Blue Detection Medium (Invivogen). TLR5 activation can then be measured by colorimetry at OD 655 nm. The cell line was cultured at 37 °C and 5% CO_2_ and maintained in DMEM media with 10% FBS and 100 U ml^−1^ of penicillin and streptomycin (Thermo Fisher Scientific), and 100 µg/mL Zeocin. The TLR5 activation assays were performed as follows: Bacterial strains were cultivated for about 2 h with agitation in LB supplemented with the appropriate antibiotics. When all the strains reached OD600nm ∼ 0.3-0.4, they were all normalized to OD = 0.3. Aliquots of the OD-normalized culture were serially diluted and plated for CFU measurements. Other aliquots of the OD-normalized culture were diluted 100 times in PBS and 20 μL of resuspended bacterial cells were pipetted into a 96-well plate. 180 μL of HEK293T-Blue-hTLR5 resuspended in HEK-Blue Detection Medium were then added on top of the agonists in each well of the plate (1.4 × 10^5^ cells/mL, corresponding to 25000 cells per well and a MOI∼20-30). Plates were incubated for 16h to 24h at 37°C, 5% CO_2_. In each well, the activation of NF-κB through TLR5 was assessed by absorbance measurements at 630 nm. Signal was then normalized according to CFU/mL and expressed as a % from the reference as indicated in each graph.

### Mice gut colonization experiment

Female C57BL/6 mice (Jackson Laboratories) were conventionally raised at Columbia University Irving Medical Center, in a rodent pathogen-free environment and in accordance with IACUC guidelines. Two separate cohorts of 5 mice were used in this study, one gavaged with *Ent Δint* (deletion of FRφ integrase to ensure the prophage cannot get excised and lost during the assay) and the other gavaged with *Ent* ΔFRφ. Mice were gavaged on three consecutive days (24h between each gavage). For the gavage step, each strain was cultivated in 150 ml LB to OD ∼ 0.3 and then renormalized to OD = 0.3. Cultures were centrifuged at 4000 *g* for 15 min and resuspended in 20 ml cold PBS in a 50 ml conical tube. They were centrifuged again at 4000 *g* for 15 minutes and resuspended in 2 ml cold PBS. The resulting suspension was washed once more in cold PBS and an aliquot was taken to assess CFU/mL (∼ 3 × 10^10^ CFU/mL averaged over the three gavage steps). The remaining cell suspension was used to gavage the mice, with 200 µL (∼ 6 × 10^9^ CFU/mL) of each strain administered to each mouse in the corresponding cohort. Faeces from each mouse in each cohort were collected (∼50 mg) before the first gavage, after the first gavage and after the last gavage at the following times: 8 hours, 1 day, 2 days, 3 days, 4 days, 5 days, 7 days, 10 days, 13 days and 25 days. Feces were weighed and resuspended in 200 µL PBS. Resuspended feces were spun down for 2 sec to pellet large dietary debris and the supernatant was serially diluted for plating. 100 µL of resuspended feces were plated on LB-Agar plates containing the appropriate antibiotic as well as serially diluted samples (ranging from 10^−1^ to 10^−4^) for each mouse of each cohort. Plates were incubated at 37 °C overnight and CFU were counted. CFU count was converted to CFU/mL and normalized by the corresponding feces mass resuspended in LB for each mouse of each cohort (CFU/(mL·mg)).

### Bioinformatic analyses of the FRφ and *Citrobacter* prophage tail fiber shufflons

To assess the recombination frequencies of FRφ tail fiber genes, high-throughput DNA sequencing of three FRφ samples was conducted by Plasmidsaurus. One sample was genomic DNA extracted from *Ent* encoding FRφ with a synthetic *cmR* insertion. Another sample was also from isolated *Ent* genomic DNA, with a *Δgin* mutation in FRφ. The third sample was genomic DNA isolated from wild type FRφ virions. FASTQ files obtained from Plasmidsaurus were imported into R, and the Biostrings package was used to probe for DNA sequencing reads that contained 25 bp of sequence at the 3′-end of the N-terminal region of the tail fiber gene concatenated to 25 bp of sequence from the 5′-end of each C-terminal region. Raw read counts corresponding to each species of tail fiber protein were graphed as a percentage of the total number of tail fiber reads using the ggplot2 package.

To assess recombination frequencies of the tail fiber shufflon from *Citrobacter amalonaticus*, three DNA sequencing datasets were downloaded from NCBI’s Short Read Archive database (SRR9220577, SRR9219931, and SRR9220572). The reads were pooled and used to construct a BLASTn database^81^, which was then queried with the sequence of the tail fiber gene that was suspected to encode the N-terminus region (WP_192927590.1 from NZ_WWUN01000014.1). A single boundary demarcated the end of the N-terminal region, beyond which different C-terminal tail fiber genes were present. Reads that didn’t span this junction, or that lacked more than 15 nt of the C-terminal region, were filtered out. The remaining reads were then manually inspected to determine which species of C-terminal tail fiber gene was present, and raw read counts were graphed using the ggplot2 package.

### Phylogenetic analysis of *csrA*-associated TldR loci

To identify *csrA*-associated TldRs and their most closely related TnpB relatives, seven sequences previously reported to be *csrA*-associated TldRs (USF27889.1, WP_251316131.1, WP_204885655.1, WP_016321625.1, WP_016324391.1, WP_087343418.1, and WP_122790109.1) were used as queries for an online BLASTp search of the NCBI NR database [default parameters]. The top 100 hits from each BLAST search were pooled, resulting in 147 unique TldR sequences which were used to again query the NCBI NR database with BLASTp [-evalue 1e-50 -qcov_hsp_perc 80 -max_hsps 50 -max_target_seqs 500]. The resulting hits were filtered to remove TnpB/TldR sequences encoded within 20 kbp of a contig end. Sequences were also clustered at an 80% amino acid identity threshold using CD-HIT^82^, before cluster representatives were aligned with MAFFT^83^ (LINSI option). A tree was then built from the resulting alignment with FastTree^84^ (-wag -gamma options) and visualized with ITOL^85^. Nuclease active site intactness was assessed by first identifying active site residues in one of the TnpB homologs (WP_087343418.1) via comparison to the ISDra2 TnpB sequence, then inspecting the multiple sequence alignment for conservation at the positions corresponding to those active site residues.

To assess genetic associations, loci were extracted from genomes encoding TnpB/TldR—comprising 20 kbp of sequence flanking each *tnpB*/*tldR* gene. These loci were then annotated with Emapper from EggNog^86^, and open reading frames (ORFs) annotated as *csrA* were extracted and translated. The CsrA ORFs were then searched against all the ORFs predicted in the *tnpB*/*tldR* loci via BLAST^81^, to ensure that none were missed or misannotated. A similar process was carried out with ORFs annotated as Flagellin, which are referred to as *fliC* for consistency, but are also denoted as *hag* in Firmicutes. TnpB/TldR homologs encoded within 3 kbp of *csrA* or *fliC* were annotated as genetically associated with those genes.

### Phylogenetic analysis of Gin proteins

To identify homologs of the *Ent* Gin protein, Gin was used to query an online BLASTp search of the NCBI Clustered_nr database [default parameters]. The top 1000 hits from the BLAST search were clustered at a 90% identity threshold using CD-HIT^82^, and then filtered to only include loci proteins encoded within loci that were at least 4-kbp length. The Gin protein sequences were then aligned with MAFFT^83^, and a tree was built from the resulting alignment using FastTree^84^ (-wag -gamma options), which was visualized in ITOL^85^. Tn3 associations were assessed via tBLASTn^81^ searches of *gin* loci, and by searching ORFs predicted in *gin* loci [getorf function of EMBOSS^87^; -table 1] with a profile HMM built from Tn3-like transposase sequences (PFAM: PF01526).

## REFERENCES

1. Mata, J., Marguerat, S. & Bähler, J. Post-transcriptional control of gene expression: a genome-wide perspective. Trends Biochem. Sci. 30, 506–514 (2005).

2. Fuda, N. J., Ardehali, M. B. & Lis, J. T. Defining mechanisms that regulate RNA polymerase II transcription in vivo. Nature 461, 186–192 (2009).

3. Browning, D. F. & Busby, S. J. W. The regulation of bacterial transcription initiation. Nature Reviews Microbiology 2, 57–65 (2004).

4. Lee, R. C., Feinbaum, R. L. & Ambros, V. The C. elegans heterochronic gene lin-4 encodes small RNAs with antisense complementarity to lin-14. Cell 75, 843–854 (1993).

5. Wiegand, T. et al. TnpB homologues exapted from transposons are RNA-guided transcription factors. Nature 631, 439–448 (2024).

6. Altae-Tran, H. et al. The widespread IS200/IS605 transposon family encodes diverse programmable RNA-guided endonucleases. Science 374, 57–65 (2021).

7. Karvelis, T. et al. Transposon-associated TnpB is a programmable RNA-guided DNA endonuclease. Nature 599, 692–696 (2021).

8. Altae-Tran, H. et al. Diversity, evolution, and classification of the RNA-guided nucleases TnpB and Cas12. Proc. Natl. Acad. Sci. 120, e2308224120 (2023).

9. Minamino, T. & Kinoshita, M. Structure, Assembly, and Function of Flagella Responsible for Bacterial Locomotion. EcoSal Plus 11, eesp-0011-2023 (2023).

10. Waldor, M. K. & Mekalanos, J. J. Lysogenic Conversion by a Filamentous Phage Encoding Cholera Toxin. Science 272, 1910–1914 (1996).

11. Wang, X. et al. Cryptic prophages help bacteria cope with adverse environments. Nature Communications 1, (2010).

12. Mirold, S. et al. Isolation of a temperate bacteriophage encoding the type III effector protein SopE from an epidemic Salmonella typhimurium strain. Proceedings of the National Academy of Sciences 96, 9845–9850 (1999).

13. Liao, H. et al. Prophage-encoded antibiotic resistance genes are enriched in human-impacted environments. Nature Communications 15, (2024).

14. Bondy-Denomy, J. et al. Prophages mediate defense against phage infection through diverse mechanisms. ISME J. 10, 2854–2866 (2016).

15. Sheahan, M. L. et al. A ubiquitous mobile genetic element changes the antagonistic weaponry of a human gut symbiont. Science 386, 414–420 (2024).

16. Wang, J. Y. & Doudna, J. A. CRISPR technology: A decade of genome editing is only the beginning. Science 379, eadd8643 (2023).

17. Nishimasu, H. et al. Crystal Structure of Cas9 in Complex with Guide RNA and Target DNA. Cell 156, 935–949 (2014).

18. Swarts, D. C., Oost, J. van der & Jinek, M. Structural Basis for Guide RNA Processing and Seed-Dependent DNA Targeting by CRISPR-Cas12a. Mol. Cell 66, 221–233.e4 (2017).

19. He, S. & Scheres, S. H. W. Helical reconstruction in RELION. Journal of Structural Biology 198, 163–176 (2017).

20. Mondino, S., Martin, F. S. & Buschiazzo, A. 3D cryo-EM imaging of bacterial flagella: Novel structural and mechanistic insights into cell motility. Journal of Biological Chemistry 298, 102105 (2022).

21. Jurczak-Kurek, A. et al. Biodiversity of bacteriophages: morphological and biological properties of a large group of phages isolated from urban sewage. Sci. Rep. 6, 34338 (2016).

22. Meynell, E. W. A phage, øχ, which attacks motile bacteria. Microbiology 25, 253–290 (1961).

23. Griffin, B. A., Adams, S. R. & Tsien, R. Y. Specific Covalent Labeling of Recombinant Protein Molecules Inside Live Cells. Science 281, 269–272 (1998).

24. Fairhead, M. & Howarth, M. Site-Specific Biotinylation of Purified Proteins Using BirA. Methods Mol. Biol. 1266, 171–184 (2014).

25. Braun, V. FhuA (TonA), the Career of a Protein. J. Bacteriol. 191, 3431–3436 (2009).

26. Quinones-Olvera, N. et al. Diverse and abundant phages exploit conjugative plasmids. Nat. Commun. 15, 3197 (2024).

27. Hayashi, F. et al. The innate immune response to bacterial flagellin is mediated by Toll-like receptor 5. Nature 410, 1099–1103 (2001).

28. Zhao, Y. et al. The NLRC4 inflammasome receptors for bacterial flagellin and type III secretion apparatus. Nature 477, 596–600 (2011).

29. Yoon, S. et al. Structural Basis of TLR5-Flagellin Recognition and Signaling. Science 335, 859–864 (2012).

30. Clasen, S. J., et al. Silent recognition of flagellins from human gut commensal bacteria by Toll-like receptor 5. Sci. Immunol. 8, eabq7001 (2023).

31. Romeo, T. & Babitzke, P. Global Regulation by CsrA and Its RNA Antagonists. Microbiol. Spectr. 6, 10.1128/microbiolspec.rwr-0009–2017 (2018).

32. Meers, C. et al. Transposon-encoded nucleases use guide RNAs to promote their selfish spread. Nature 622, 863–871 (2023).

33. Drider, D. & Condon, C. The Continuing Story of Endoribonuclease III. J. Mol. Microbiol. Biotechnol. 8, 195–200 (2004).

34. Nashimoto, H. & Uchida, H. DNA sequencing of the Escherichia coli ribonuclease III gene and its mutations. Mol. Gen. Genet. MGG 201, 25–29 (1985).

35. Deltcheva, E. et al. CRISPR RNA maturation by trans-encoded small RNA and host factor RNase III. Nature 471, 602–607 (2011).

36. Shmakov, S. et al. Discovery and Functional Characterization of Diverse Class 2 CRISPR-Cas Systems. Mol. Cell 60, 385–397 (2015).

37. Johnson & Reid, C. Site-specific DNA Inversion by Serine Recombinases. Microbiology Spectrum 3, (2015).

38. Silverman, M., Zieg, J., Hilmen, M. & Simon, M. Phase variation in Salmonella: genetic analysis of a recombinational switch. Proc. Natl. Acad. Sci. 76, 391–395 (1979).

39. Zetsche, B. et al. Cpf1 Is a Single RNA-Guided Endonuclease of a Class 2 CRISPR-Cas System. Cell 163, 759–771 (2015).

40. Klompe, S. E., Vo, P. L. H., Halpin-Healy, T. S. & Sternberg, S. H. Transposon-encoded CRISPR– Cas systems direct RNA-guided DNA integration. Nature 571, 219–225 (2019).

41. Strecker, J. et al. RNA-guided DNA insertion with CRISPR-associated transposases. Science 365, 48–53 (2019).

42. Qi, L. S. et al. Repurposing CRISPR as an RNA-Guided Platform for Sequence-Specific Control of Gene Expression. Cell 152, 1173–1183 (2013).

43. Heller & Knut, J. Molecular interaction between bacteriophage and the gram-negative cell envelope. Archives of Microbiology 158, 235–248 (1992).

44. Taylor, V. L. et al. Prophages block cell surface receptors to ensure survival of their viral progeny. (2024) doi:10.1101/2024.03.05.583538.

45. Sztanko, K. M. et al. Prophages express a type IV pilus component to provide anti-phage defence. (2024) doi:10.1101/2024.03.29.587342.

46. Lisevich, I., Colin, R., Yang, H. Y., Ni, B. & Sourjik, V. Physics of swimming and its fitness cost determine strategies of bacterial investment in flagellar motility. Nature Communications 16, (2025).

47. Bardoel, B. W. et al. Pseudomonas Evades Immune Recognition of Flagellin in Both Mammals and Plants. PLoS Pathogens 7, e1002206 (2011).

48. Ribet, D. & Cossart, P. How bacterial pathogens colonize their hosts and invade deeper tissues. Microbes and Infection 17, 173–183 (2015).

49. Josenhans, C. & Suerbaum, S. The role of motility as a virulence factor in bacteria. International Journal of Medical Microbiology 291, 605–614 (2002).

50. Arthur, T. D. et al. Invertible promoters mediate bacterial phase variation, antibiotic resistance, and host adaptation in the gut. Science 363, 181–187 (2019).

51. Klose, K. E. & Mekalanos, J. J. Differential Regulation of Multiple Flagellins inVibrio cholerae. J. Bacteriol. 180, 303–316 (1998).

52. Klose, K. E. & Mekalanos, J. J. Differential Regulation of Multiple Flagellins in Vibrio cholerae. J. Bacteriol. 180, 303–316 (1998).

53. Sharan, S. K., Thomason, L. C., Kuznetsov, S. G. & Court, D. L. Recombineering: a homologous recombination-based method of genetic engineering. Nat. Protoc. 4, 206–223 (2009).

54. Reisch, C. R. & Prather, K. L. J. Scarless Cas9 Assisted Recombineering (no-SCAR) in Escherichia coli, an Easy-to-Use System for Genome Editing. Curr. Protoc. Mol. Biology 117, 31.8.1–31.8.20 (2017).

55. Palma, V., Gutiérrez, M. S., Vargas, O., Parthasarathy, R. & Navarrete, P. Methods to Evaluate Bacterial Motility and Its Role in Bacterial–Host Interactions. Microorganisms 10, 563 (2022).

56. Kreutzberger, M. A. B. et al. Flagellin outer domain dimerization modulates motility in pathogenic and soil bacteria from viscous environments. Nat. Commun. 13, 1422 (2022).

57. Rohou, A. & Grigorieff, N. CTFFIND4: Fast and accurate defocus estimation from electron micrographs. J. Struct. Biol. 192, 216–221 (2015).

58. Lövestam, S. & Scheres, S. H. W. High-throughput cryo-EM structure determination of amyloids. Faraday Discussions 240, 243–260 (2022).

59. Bepler, T. et al. Positive-unlabeled convolutional neural networks for particle picking in cryo-electron micrographs. Nat. Methods 16, 1153–1160 (2019).

60. Egelman & Edward, H. A robust algorithm for the reconstruction of helical filaments using single-particle methods. Ultramicroscopy 85, 225–234 (2000).

61. Trachtenberg, S., DeRosier, D. J., Aizawa, S.-I. & Macnab, R. M. Pairwise perturbation of flagellin subunits. Journal of Molecular Biology 190, 569–576 (1986).

62. Zivanov, J., Nakane, T. & Scheres, S. H. W. A Bayesian approach to beam-induced motion correction in cryo-EM single-particle analysis. IUCrJ 6, 5–17 (2019).

63. Pettersen, E. F. et al. UCSF Chimera—A visualization system for exploratory research and analysis. Journal of Computational Chemistry 25, 1605–1612 (2004).

64. Afonine, P. V. et al. Real-space refinement inPHENIXfor cryo-EM and crystallography. Acta Crystallographica Section D Structural Biology 74, 531–544 (2018).

65. Murshudov, G. N. et al. REFMAC5 for the refinement of macromolecular crystal structures. Acta Crystallographica Section D Biological Crystallography 67, 355–367 (2011).

66. Nicholls, R. A., Fischer, M., McNicholas, S. & Murshudov, G. N. Conformation-independent structural comparison of macromolecules withProSMART. Acta Crystallographica Section D Biological Crystallography 70, 2487–2499 (2014).

67. Zhao, Z. et al. Frequent pauses in Escherichia coli flagella elongation revealed by single cell real-time fluorescence imaging. Nat. Commun. 9, 1885 (2018).

68. Schneider, C. A., Rasband, W. S. & Eliceiri, K. W. NIH Image to ImageJ: 25 years of image analysis. Nat. Methods 9, 671–675 (2012).

69. Bonocora, R. P. & Wade, J. T. ChIP-Seq for Genome-Scale Analysis of Bacterial DNA-Binding Proteins. Methods Mol. Biol. 1276, 327–340 (2015).

70. Chen, S., Zhou, Y., Chen, Y. & Gu, J. fastp: an ultra-fast all-in-one FASTQ preprocessor. Bioinformatics 34, i884–i890 (2018).

71. Langmead, B. & Salzberg, S. L. Fast gapped-read alignment with Bowtie 2. Nat. Methods 9, 357–359 (2012).

72. Danecek, P. et al. Twelve years of SAMtools and BCFtools. GigaScience 10, giab008 (2021).

73. Ramírez, F., et al. deepTools2: a next generation web server for deep-sequencing data analysis. Nucleic Acids Res. 44, W160–W165 (2016).

74. Li, H. & Durbin, R. Fast and accurate short read alignment with Burrows–Wheeler transform. Bioinformatics 25, 1754–1760 (2009).

75. Martin, M. Cutadapt removes adapter sequences from high-throughput sequencing reads. EMBnetJ. 17, 10 (2011).

76. Robinson, J. T. et al. Integrative genomics viewer. Nat. Biotechnol. 29, 24–26 (2011).

77. Banks, D. J., Lei, B. & Musser, J. M. Prophage Induction and Expression of Prophage-EncodedVirulence Factors in Group A Streptococcus Serotype M3 StrainMGAS315. Infect. Immun. 71, 7079–7086 (2003).

78. Tang, S. et al. De novo gene synthesis by an antiviral reverse transcriptase. Science 386, eadq0876 (2024).

79. Li, N. et al. Characterization of Phage Resistance and Their Impacts on Bacterial Fitness in Pseudomonas aeruginosa. Microbiol. Spectr. 10, e02072–22 (2022).

80. Kille, B. et al. Parsnp 2.0: scalable core-genome alignment for massive microbial datasets. Bioinformatics 40, btae311 (2024).

81. Camacho, C. et al. BLAST+: architecture and applications. BMC Bioinform. 10, 421 (2009).

82. Huang, Y., Niu, B., Gao, Y., Fu, L. & Li, W. CD-HIT Suite: a web server for clustering and comparing biological sequences. Bioinformatics 26, 680–682 (2010).

83. Katoh, K. & Standley, D. M. MAFFT Multiple Sequence Alignment Software Version 7: Improvements in Performance and Usability. Mol. Biol. Evol. 30, 772–780 (2013).

84. Price, M. N., Dehal, P. S. & Arkin, A. P. FastTree 2 – Approximately Maximum-Likelihood Trees for Large Alignments. PLoS ONE 5, e9490 (2010).

85. Letunic, I. & Bork, P. Interactive Tree Of Life (iTOL) v5: an online tool for phylogenetic tree display and annotation. Nucleic Acids Res. 49, W293–W296 (2021).

86. Cantalapiedra, C. P., Hernández-Plaza, A., Letunic, I., Bork, P. & Huerta-Cepas, J. eggNOG-mapper v2: Functional Annotation, Orthology Assignments, and Domain Prediction at the Metagenomic Scale. Mol. Biol. Evol. 38, 5825–5829 (2021).

87. Rice, P., Longden, I. & Bleasby, A. EMBOSS: The European Molecular Biology Open Software Suite. Trends Genet. 16, 276–277 (2000).

