## Extended Data for "Temperate phages enhance host fitness via RNA-guided flagellar remodeling"

### EXTENDED DATA FIGURES

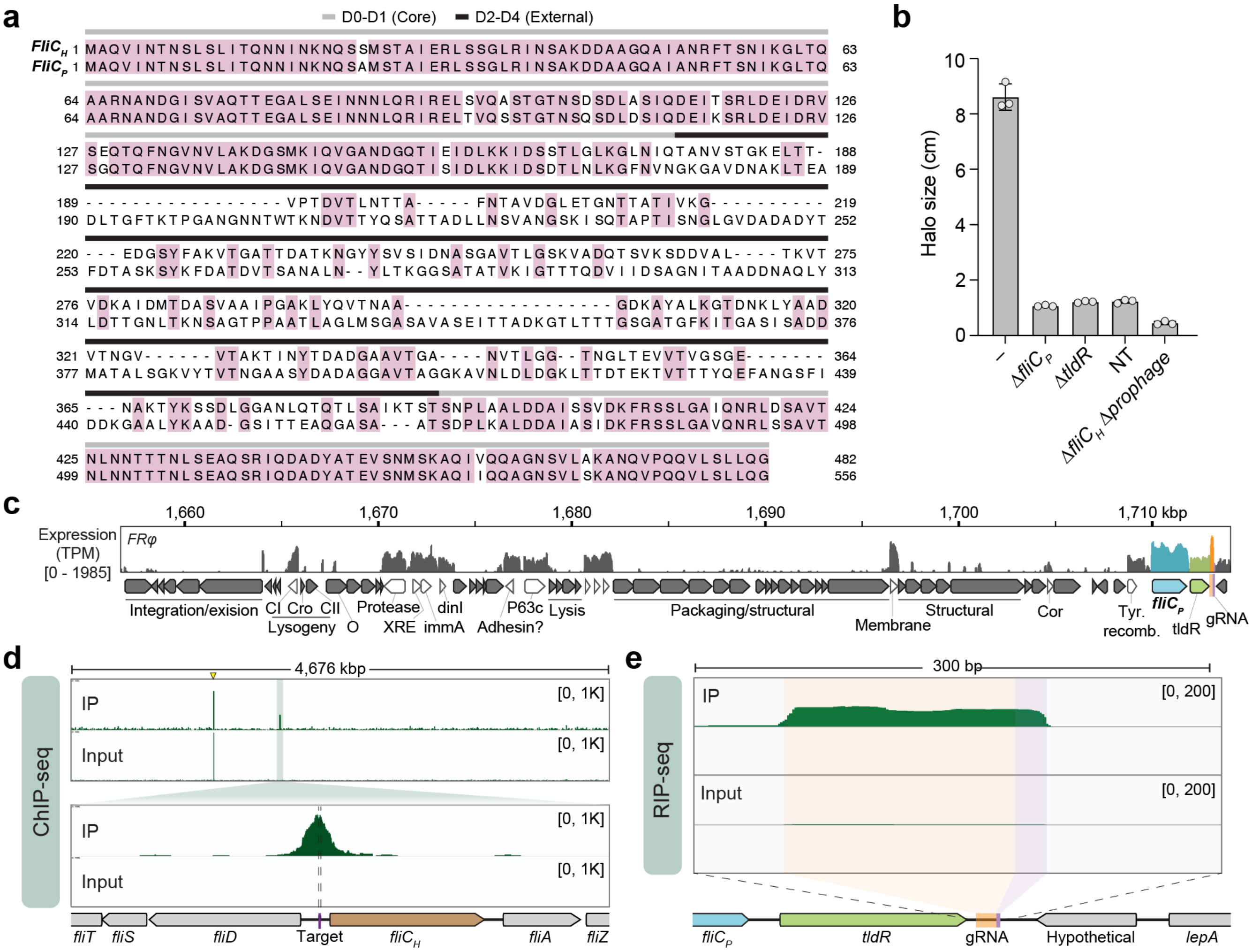


**Extended Data Fig. 1 | Native flagellin regulation by FRφ-encoding TldR: sequence alignment, expression, and targeting data.**

**(a)** *Enterobacter* sp. BIDMC93 (*Ent*) FliC$_H$ and FliC$_P$ amino acid similarity; pink coloring indicates identical residues.

**(b)** Bar graph quantifying bacterial motility in *Ent* strains with the indicated prophage mutations. Bars indicate mean ± s.d. (n = 3 biological replicates).

**(c)** Genomic architecture of FRφ alongside RNA-seq of the WT lysogenic strain (reproduced from Wiegand *et al*. 2024[11]).

**(d)** ChIP-seq coverage profiles of FLAG-tagged *Ent*TldR expressed in its native *Enterobacter* sp. BIDMC93 cellular context (top), alongside a magnified inset showing a 5-kb window surrounding the host *fliC$_H$* locus (bottom). The prominent peak in the top panel (yellow arrowhead) corresponds to the *tldR* locus and reflects increased signal in both IP and Input due to plasmid-based overexpression of the FLAG-TldR used for ChIP-seq.

**(e)** RIP-seq coverage profiles of FLAG-tagged *Ent*TldR expressed in its native *Enterobacter* sp. BIDMC93 cellular context, showing native association with the guide RNA. The guide sequence is highlighted in purple, and the upstream scaffold is shown in orange.

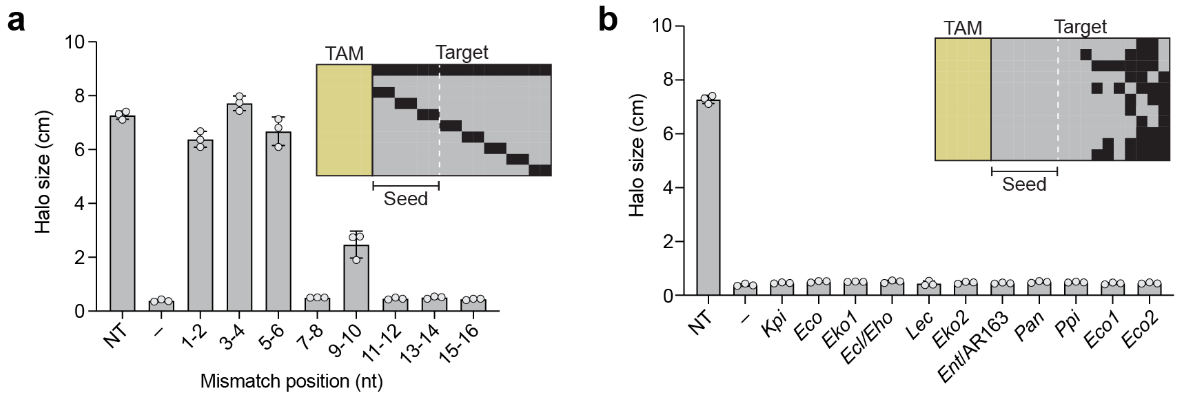


**Extended Data Fig. 2 | Guide RNA specificity determines TldR-mediated repression of bacterial motility**

**(a)** Bar graph quantifying bacterial motility in *E. coli* AW405 strains expressing TldR and guide RNAs containing 2-nt mismatches, assessing guide RNA sequence requirements for *fliC* repression. NT, non-targeting; "–", WT (fully matching) guide. The schematic above illustrates the positions of specific mutations (black rectangles) relative to an invariant cognate target adjacent motif (TAM). Bars indicate mean ± s.d. (n = 3 biological replicates).

**(b)** Bar graph quantifying bacterial motility in *E. coli* AW405 strains expressing TldR and guide RNAs with native mismatches, assessing guide RNA sequence requirements for *fliC* repression. NT, non-targeting; "–", WT (fully matching) guide. The schematic above illustrates the positions of specific mutations (black rectangles) relative to an invariant cognate TAM. Bars indicate mean ± s.d. (n = 3 biological replicates).

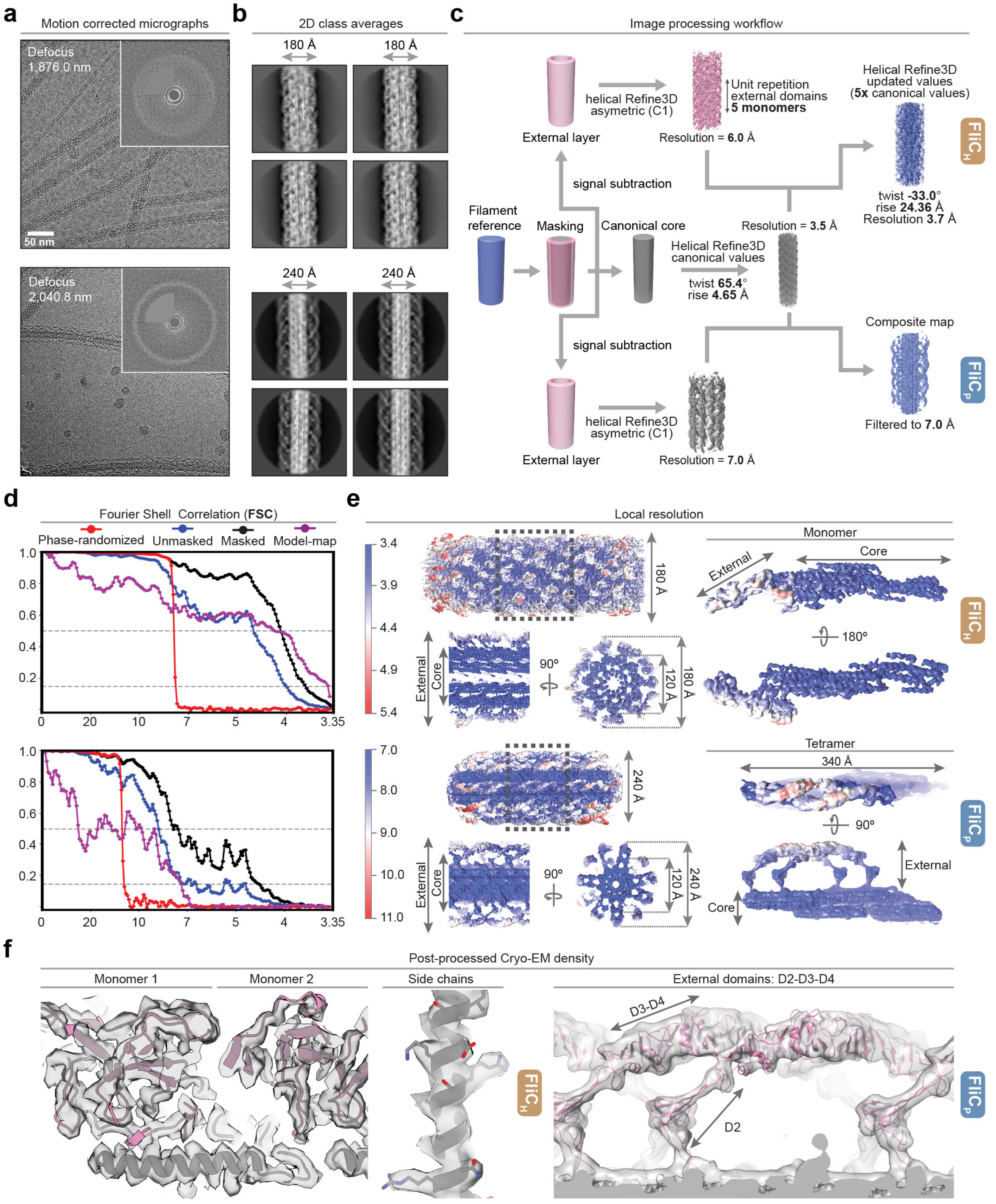


**Extended Data Fig. 3 | Cryo-EM data collection, refinement, and structure determination.**

(**a**) Representative motion corrected cryo-EM micrographs for samples of FliC$_H$ (top) and FliC$_P$ (bottom).

(**b**) Representative 2D class averages reconstructed from FliC$_H$ (top) and FliC$_P$ (bottom) micrographs.

(**c**) Schematic representation of the image processing workflow followed for the helical reconstruction of FliC$_H$ and FliC$_P$ filaments.

(**d**) Fourier shell correlation (FSC) curves for the final reconstructed maps and final refined models.

(**e**) Local resolution maps of FliC$_H$ (top) and FliC$_P$ (bottom) reconstructed filaments.

(**f**) Post-processed cryo-EM density maps in representative regions of FliC$_H$ (left) and FliC$_P$ (right) support the placement of secondary structural features and residue side chains.

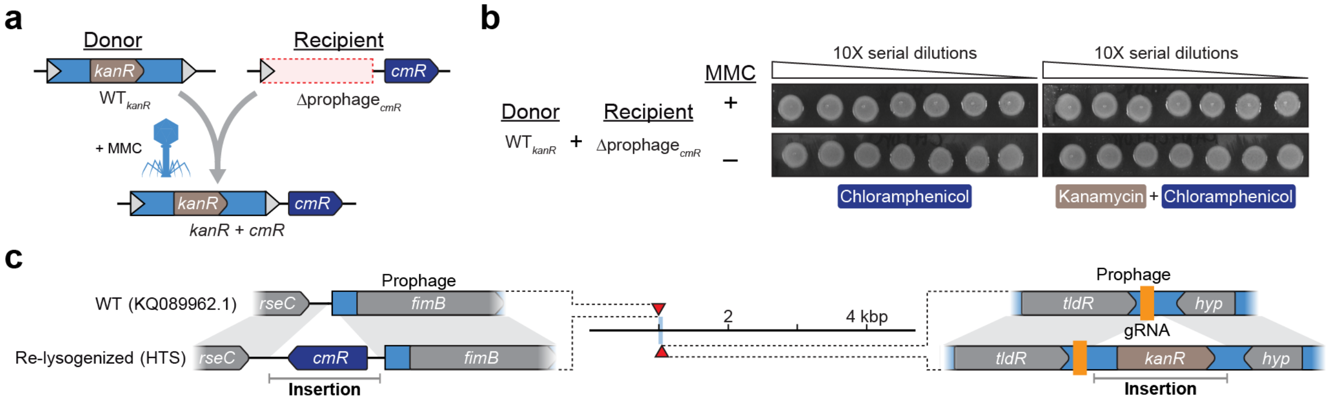


**Extended Data Fig. 4 | Controls validating the lysogenization assay using antibiotic resistant markers.**

**(a)** Schematic of lysogenization assay.

**(b)** Representative plate images after lysogenization experiments in the presence or absence of mitomycin C (MMC).

(**b**) High-throughput sequencing of genomic DNA reveals that doubly drug-resistant clones harbor genomic *cmR* insertions (left) and prophage-encoded *kanR* genes (right) relative to the WT genome, indicative of novel lysogenization events.

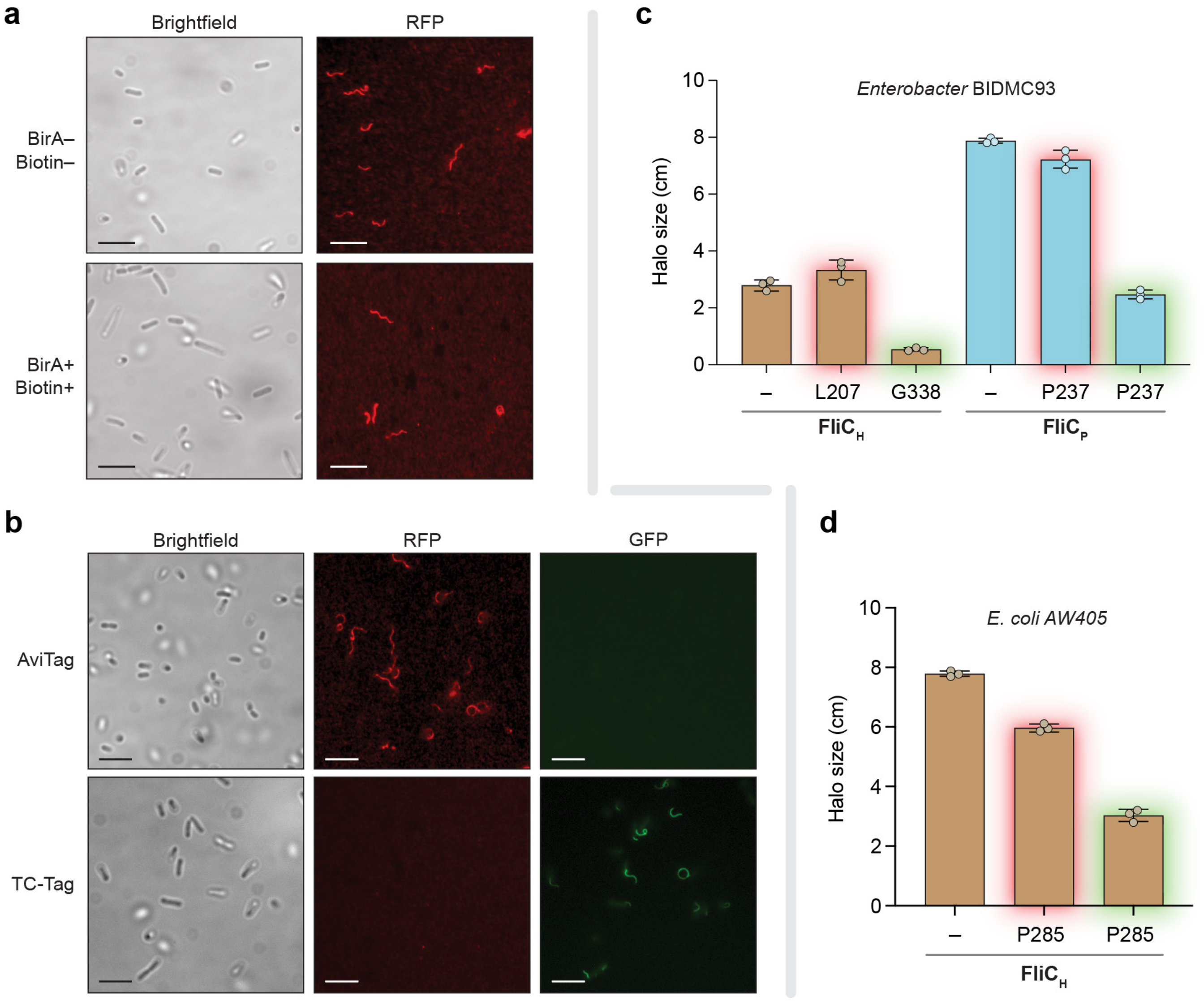


**Extended Data Fig. 5 | Validation of orthogonal flagellin labeling and functional motility of tagged constructs.**

**(a)** Representative microscopy images of *E. coli* MG1655 Δ*fliC* cells with AviTagged FliC in the absence or presence of exogenously added BirA and biotin, demonstrating that biotin and BirA are not required for flagellin labeling and detection with streptavidin-Alexa Fluor 568. Scale bar represents 5 µm.

**(b)** Representative microscopy images of *E. coli* MG1655 AviTag (top) and TC-tag (bottom) FliC cells that have each been co-labeled by both FlAsH and streptavidin-Alexa Fluor 568, demonstrating that the two labeling approaches exclusively label their respective tagged flagellin. Scale bar represents 5 µm.

**(c)** Bar graph quantifying bacterial motility in *Ent* strains encoding endogenously tagged versions of either $fliC_H$ or $fliC_P$. Tag type is indicated by the bar halo color (AviTag in red, TC-tag in green), and tag position is shown on the X-axis. $fliC_H$ strains are in a Δ*prophage* background, and $fliC_P$ strains are in a $\Delta fliC_H$ background; "–" indicates untagged strains. Bars indicate mean ± s.d. (n = 3 biological replicates).

**(d)** Bar graph quantifying bacterial motility in *E. coli* AW405 strains encoding *fliC* endogenously tagged with either AviTag (red) or TC-tag (green) at position P285. Plates were imaged for halo-size measurements after 16 hours of growth. Bars indicate mean ± s.d. (n = 3 biological replicates).

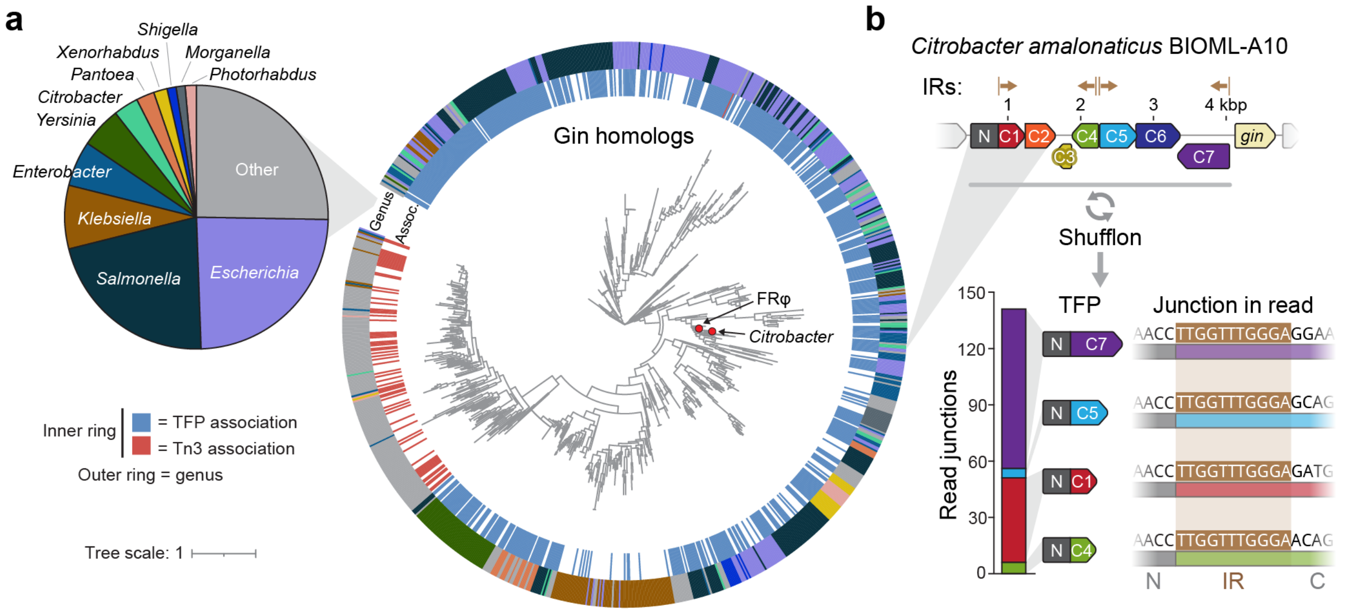


**Extended Data Fig. 6 | Phylogenetic insights into Gin-associated shufflon loci encoding tail fiber protein diversity**

**(a)** Phylogenetic tree built on Gin protein sequences showing genetic associations with the tail fiber protein (TFP, blue) or Tn3 transposase (red, inner ring), revealing a homologous relationship to Tn3 invertase-family proteins (TnpR). The pie chart (left) indicates the abundance of each genus represented in the tree (outer ring).

(**b**) Shufflon from a prophage in *Citrobacter amalonaticus* [NCBI: NZ_WWUN01000014.1] that encodes tail fiber protein (TFP) isoforms with an invariable N-terminus and seven C-terminal variants (top). DNA sequencing reads from three SRA datasets (SRR9220577, SRR9219931, SRR9220572) indicate that at least four tail fiber species are regularly sampled, with junctions between the N- and C-termini demarcated by a conserved inverted repeated (IR) that is likely recognized by Gin (bottom).

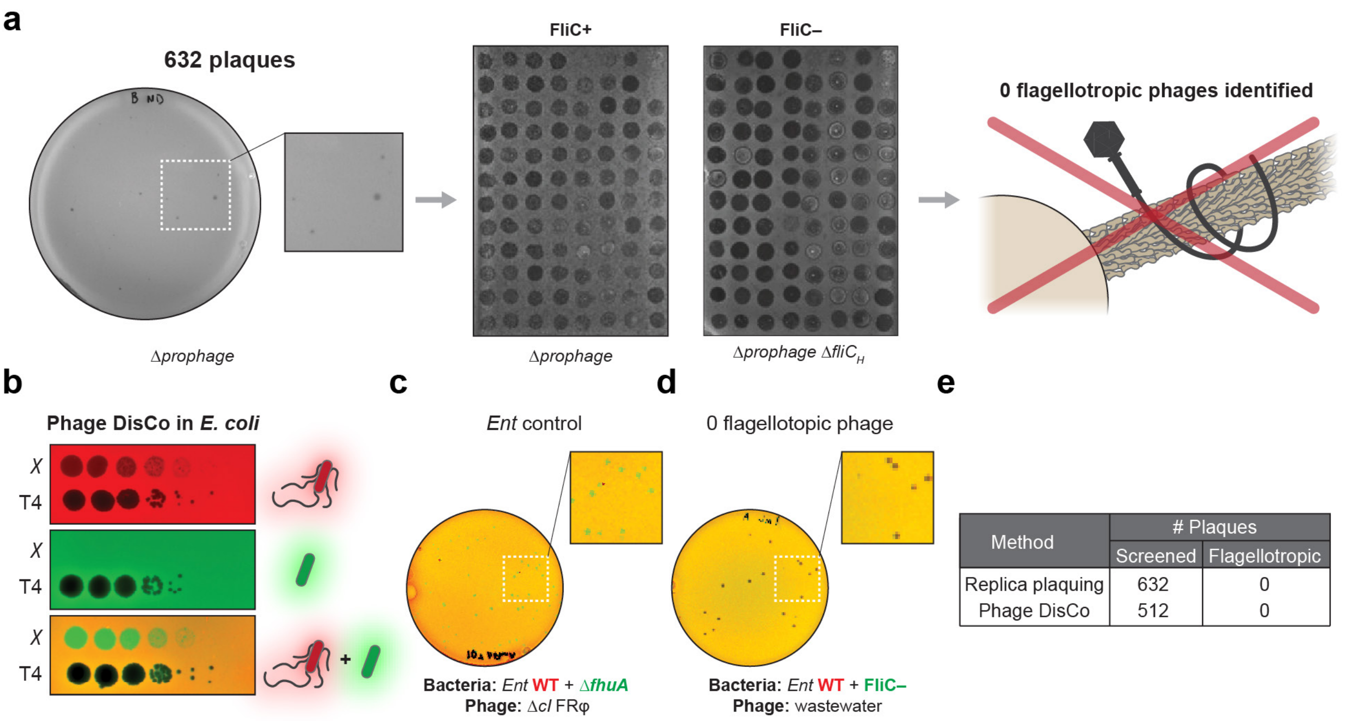


| Method | # Plaques | |
|---|---|---|
| | Screened | Flagellotropic |
| Replica plaquing | 632 | 0 |
| Phage DisCo | 512 | 0 |

**Extended Data Fig. 7 | Wastewater screening strategies reveal no evidence of flagellotropic phages infecting *Enterobacter*.**

(**a**) Screening for flagellotropic phages in wastewater using a replica plaquing strategy. The left image shows a representative plaque assay using Δ*prophage Ent* cells to isolate phages from environmental samples, which were then replica-plated on parallel strains either lacking or expressing *fliC*$_H$ (middle panels). No phages were identified with this strategy that selectively infected flagellated cells but not non-flagellated cells, suggesting an absence of flagellotropic phages in the screened samples.

**(b)** Phage DisCo controls to identify flagellotropic phages in *E. coli* (spot assay), using both χ phage (flagellotropic) and T4 phage (non-flagellotropic). The top two images are plaque assays using monocultures of WT *E. coli* with red fluorescence (top) or Δ*fliC E. coli* with green fluorescence (middle); the bottom image is of a co-culture, where both strains were mixed before plaquing.

**(c)** Representative plate from a Phage DisCo control assay using FRφ Δ*cI* phage plated on a mixed lawn of WT and Δ*fhuA Ent* strains labeled with red and green fluorescence, respectively. Green fluorescent plaques indicate regions where the Δ*cI* phage has lysed red WT cells but failed to infect the green Δ*fhuA* cells, consistent with an *fhuA*-dependent infection mechanism.

**(d)** Representative plate from a Phage DisCo screen of environmental samples for flagellotropic phages, using a mixed lawn of WT and FliC– (Δ*prophage* Δ*fliC*$_H$) *Ent* strains labeled with red and green fluorescence, respectively. No fluorescent plaques were observed, indicating that none of the isolated phages selectively infected the flagellated WT strain.

**(e)** Table summarizing the number of plaques screened using each approach.

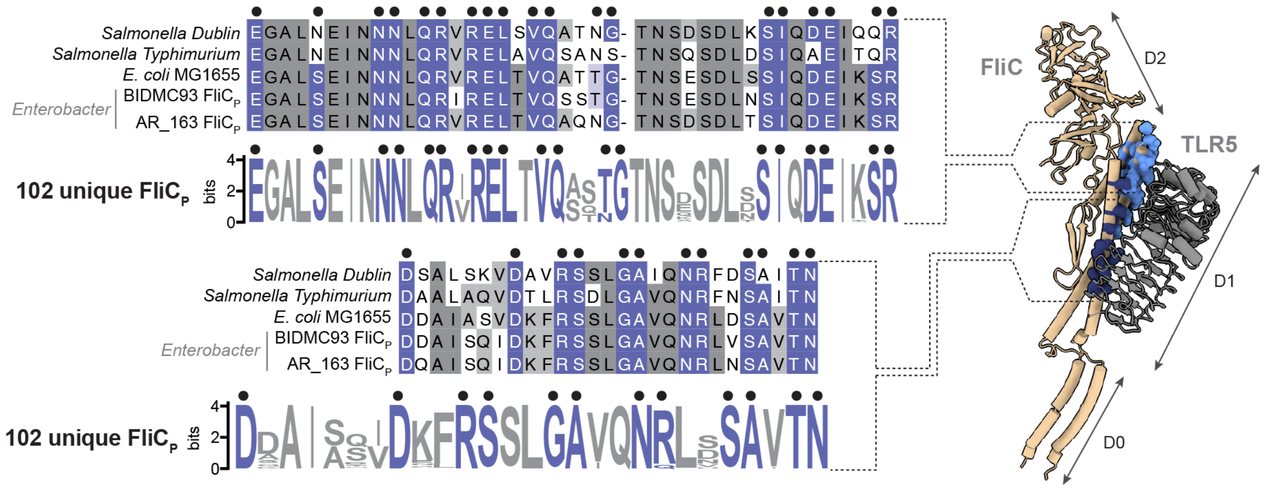


**Extended Data Fig. 8 | Comparison of FliC epitopes recognized by TLR5.**

Multiple sequence alignment (MSA) of the indicated FliC homologs (left), focusing on the motifs recognized by TLR5 in two different regions of the D1 domain (contacts denoted by black dots). Conserved residues that are, or are not, part of the epitope are highlighted in blue and grey, respectively. Weblogos built from corresponding regions of a broader $FliC_P$ alignment (n = 102 unique sequences) are shown below each MSA. Epitopes are highlighted on a structure of FliC from *Salmonella typhimurium* (PDB: 1UCU) that was superimposed on a flagellin bound structure of TLR5 from *Danio rerio* (PDB: 3V47) (right).

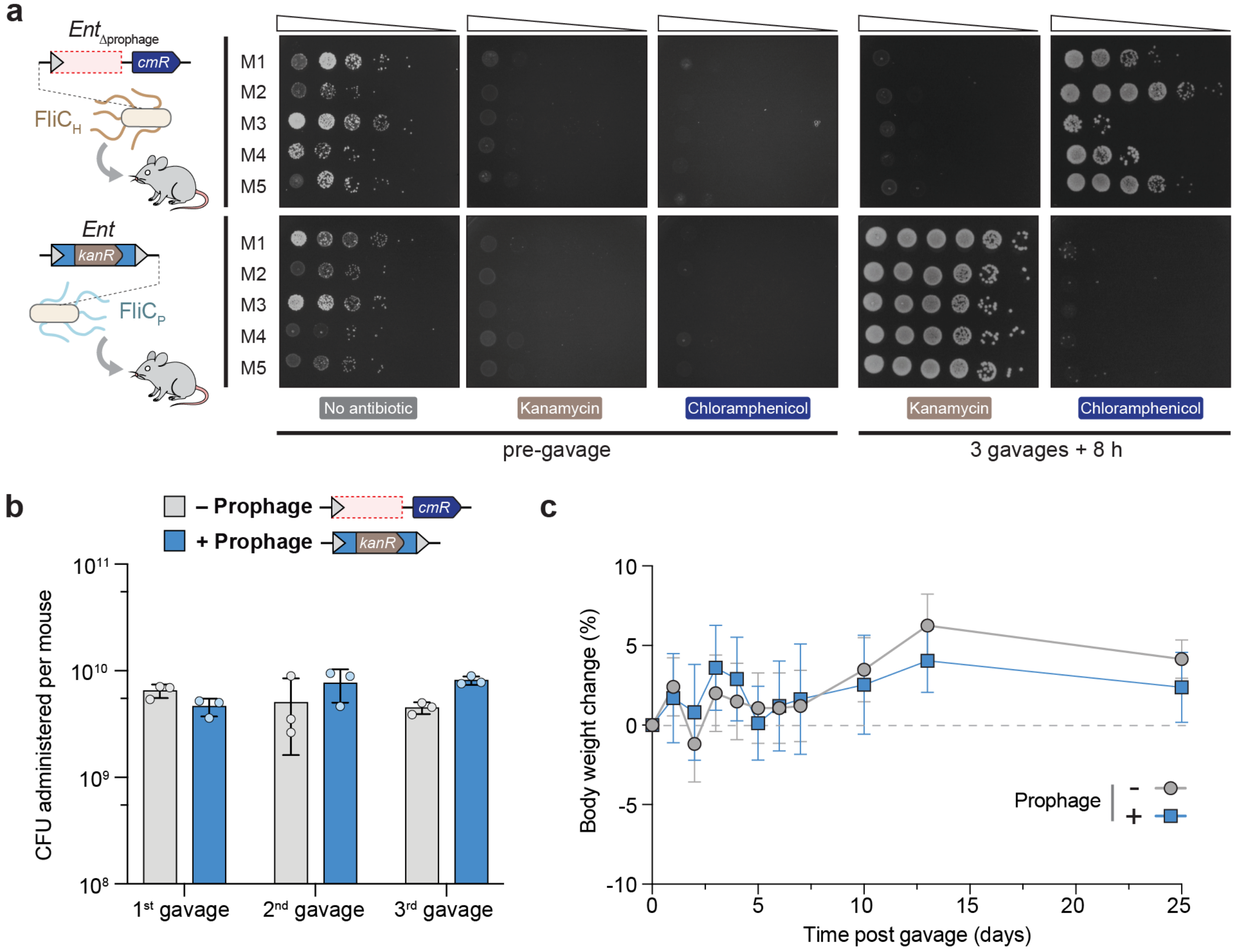


**Extended Data Fig. 9 | Characterization of bacterial colonization from *in vivo* experiments.**

**(a)** Spot assay obtained from resuspending mice feces in PBS and plating on agar plates containing the indicated antibiotic, to select for either *Ent* Δ*prophage* (CmR strain) or *Ent* +prophage (KanR strain). No CmR or KanR strains were observed prior to gavage, despite the presence of bacteria on the plate lacking antibiotic, whereas strains with the expected resistance grow on agar plates of the corresponding group 8 h after the last gavage.

**(b)** Colony-forming units (CFU) administered per mouse, as quantified by plating on each day of gavage for each group. Each individual value corresponds to a technical replicate originating from one single bacterial suspension of the corresponding strain, and 200 µl of each suspension was used to gavage each mouse from the corresponding group.

**(c)** Effect of gavaging mice with *Ent* lacking or containing the FRφ prophage on body weight, used as a readout of mouse general health.

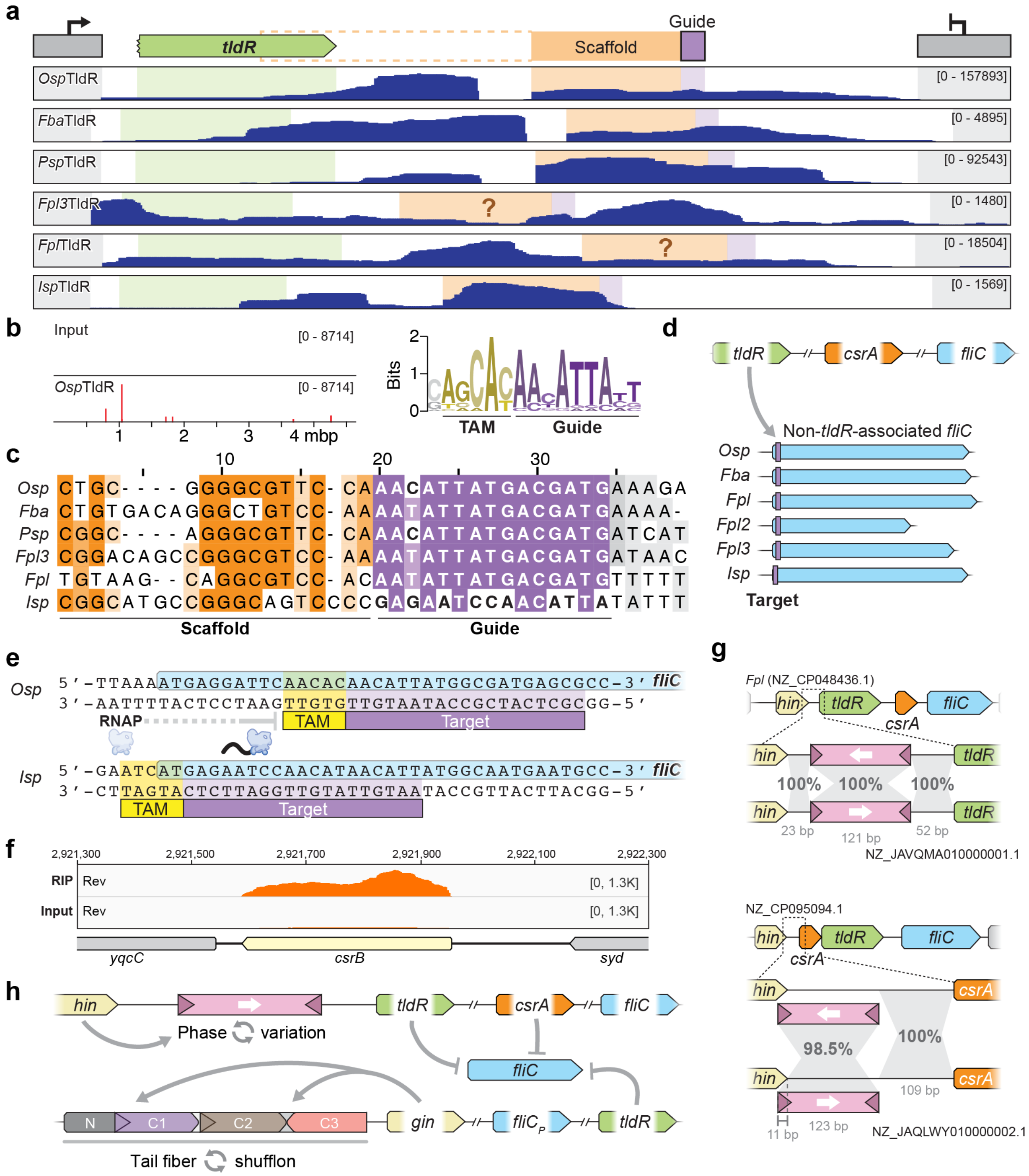


**Extended Data Fig. 10 | Guide-target identification for CsrA-associated TldR homologs.**

**(a)** RIP-seq coverage tracks of six distinct *csrA*-associated TldR homologs. The purple highlight indicates the putative gRNA guide sequence, and the orange highlight indicates the putative gRNA scaffold sequence.

**(b)** ChIP-seq coverage of the *E. coli* genome in cells expressing TldR from *Oscillibacter* sp. 1-3 (left), and WebLogo of motif extracted from peaks corresponding to the *Osp*TldR TAM and guide sequences (right).

**(c)** Sequence alignment of putative guides for the six *csrA*-associated TldR homologs shown in (a).

**(d)** Putative targets of *csrA*-associated TldR guides at the 5′ end of a *fliC* gene encoded distally in the genome, which are distinct from the corresponding sequence in *tldR*-associated *fliC* genes.

**(e)** Nucleotide-level view of the *Osp*TldR target and TAM at the 5′ end of *fliC* (top), which is representative of the positioning of targets for *Fba*, *Fpl*, *Fpl2*, and *Fpl3*TldR. The putative target of *Isp*TldR is shifted relative to these targets, and all of these targets are expected to block RNAP elongation, terminating *fliC* transcription.

(**f**) RIP-seq coverage profiles of FLAG-tagged *Osp*CsrA at the *E. coli csrB* locus.

(**g**) Comparative genomics reveals a region in the *Flavonifractor plautii* JCM 32125 genome that exhibits local inversion relative to a closely related genome (top), suggesting that the accompanying *hin* gene product governs phase variation for the downstream *tldR* gene. A similar region is inverted in two closely related *F. plautii* genomes (bottom), upstream of *csrA*. Triangles in the inverted regions represent inverted repeats, and white arrows indicate the direction of transcription based on the presence of a FliA promoter motif (PRODORIC database: MX000132).

(**h**) Convergent mechanisms of *fliC* gene regulation by *csrA*-associated and *fliC*$_P$-associated TldRs, and similar DNA inversion loci that control phase variation of *tldR* loci and phage tail fiber (TFP) proteins.
